# Single-nucleus RNA sequencing reveals enduring signatures of acute stress and chronic exercise in striatal microglia

**DOI:** 10.1101/2024.06.24.600469

**Authors:** Meghan G. Connolly, Zachary V. Johnson, Lynna Chu, Nicholas D. Johnson, Trevor J. Buhr, Elizabeth M. McNeill, Peter J. Clark, Justin S. Rhodes

## Abstract

Acute stress has enduring effects on the brain and motivated behavior across species. For example, acute stress produces persisting decreases in voluntary physical activity as well as molecular changes in the striatum, a brain region that regulates voluntary physical activity and other motivated behaviors. Microglia, the primary immune cells of the central nervous system, are positioned at the interface between neural responses to stress and neural coordination of voluntary activity in that they respond to stress, sense molecular changes in the striatum, and modulate neuronal activity. However, the role of striatal microglia in stress-induced long-term suppression of voluntary activity is unknown. Here we employ single nucleus RNA-sequencing to investigate how stress and exercise impact the biology of microglia in the striatum. We find that striatal microglia display altered activation profiles six weeks after an acute stressor. Furthermore, we show that access to a running wheel is associated with an additional and distinct microglial activation profile characterized by upregulation of genes related to complement components and phagocytosis pathways. Lastly, we find that distinct gene sets show expression changes associated with general access to a running wheel versus variation in running levels. Taken together, our results deepen our understanding of the diverse molecular states that striatal microglia assume in response to stress and exercise and suggest that microglia exhibit a broader range of functional states than previously thought.

## 1. Introduction

Adverse life events (e.g., psychological traumas or severe stressors) can have lasting consequences on human health^1,2^. Keeping physically active is crucial for maintaining physical and mental health and is considered one of the best ways of coping with stress^3,4^. However, exposure to adverse events can consequently lower physical activity engagement across the lifespan ^5,6^. A better understanding of how adverse experiences chronically impede physical activity engagement is critical for conceptualizing new approaches for improving exercise engagement across the lifespan.

Similarly to humans, rodents exposed to stressors of sufficient intensity can develop persistent reductions in physical activity engagement, offering a useful model for investigating the interactions between stress and physical inactivity. Rodents have a natural inclination to run. For example, rats will lever press for access to a running wheel, indicating that they find wheel running to be rewarding^7–10^. However, rats exposed to a single episode of inescapable tail shocks (acute stress) display a four-fold decrease in voluntary wheel running that outlasts the development of all other well-characterized anxiety– and depression-like behaviors by months^11,12^. Moreover, rats exposed to acute stress can perform forced physical activity at high levels, as performance on treadmill running to failure^11^, 15 minutes of forced swim^13^, and approximately an hour of shuttle box escape testing^14,15^ recovers to the levels of non-stressed rats within a week indicating that the ability to exercise is still present. Taken together, these data suggest that long-term reductions in voluntary wheel running following acute stress may represent a persistent deficit in motivation specifically towards physically exertive voluntary activities.

Multiple regions of the brain contribute to motivation for voluntary physical activity^16^. The striatum, both its dorsal and ventral areas, is particularly notable for its dual role in motivating reward-seeking behaviors and regulating movement^17–21^. A recent study found major transcriptomic changes in this region in mice that were selectively bred for increased voluntary wheel running behavior^22^. The striatum is also well known to be sensitive to stress, with several studies showing stress alters striatal circuitry^23^ and striatal secretion of serotonin and dopamine^24^. Additional studies have shown that stress alte rs striatal-dependent processes like motivation, locomotion, and associative learning^23,25^. Given that the striatum is both sensitive to stress and regulates reward information processing, decision-making, and motivated behaviors^21^, this brain region is a lead candidate in which the neurobiological impacts of stress may underlie stress-associated decreases in motivation for voluntary physical activity. However, the specific cell types and molecular changes in the striatum that remain impacted long after an acute stressor remain unknown.

Microglia are primely positioned to regulate striatal responses to stress and striatal-dependent motivated behaviors owing to their roles in neuroinflammation and homeostatic regulation^26–29^. In response to an acute stressor like inescapable shock, microglia enact a robust but brief neuroinflammatory response^30–32^ that subsides within approximately 48 hours. However, some microglia remain in a “primed” state for several weeks after exposure to a stressor ^33–37^. This primed state is distinctively marked by increased baseline levels of inflammation markers and inflammatory mediators, a reduced threshold for activation into a pro-inflammatory state, and an intensified inflammatory response upon activation ^33,35,37,38^. In addition to their traditional role as mediators of inflammation, microglia are increasingly recognized for their specialized functions in regulating the extracellular microenvironment in a way that influences brain-region specific signaling^39^. For instance, striatal microglia detect and react to neuronal ATP release by increasing production of adenosine, which in turn suppresses neuronal activity through adenosine receptors^40^. The responses of striatal microglia to stress may therefore contribute to long-term alterations in striatal function and motivational deficits.

However, the specific molecular profile of microglia long after an acute stressor that occurs in coincidence with prolonged reduced voluntary physical activity has not been characterized to our knowledge. Here we use single nucleus RNA-sequencing to investigate alterations in striatal microglia six weeks after acute stress in relation to prolonged decreases in motivation for physical activity. To evaluate microglial signatures of stress and exercise, male rats were given one session of inescapable shock stress and then allowed access to a running wheel for 42 days. Compared to controls, we predicted that microglia in the striatum of acutely stressed rats would show enduring and distinct forms of neuroinflammatory activation, and that physical activity would promote an anti-inflammatory microglial phenotype. Our results further our understanding of microglial functional complexity and reveal candidate molecular and cellular mechanisms underlying long-term stress-associated decreases in motivation for physical activity.

## 2. Materials and Methods

### 2.1 Subjects

Adult male Sprague Dawley (SD) rats from Envigo (250-280g) were single-housed upon arrival to the Iowa State University Vivarium. Male rats were used for this study as the vast majority of over five decades of research on stress physiology using tail shock model has been completed in this exact paradigm using male rats, which could provide important behavioral or physiological context to outcomes in this study^41–46^. Moreover, long-term running deficits following acute stress have been previously documented in male, but not yet female, rats ^11,12^. Chow (ENVIGO, Tekland 2014) and water were provided *ad libitum* throughout the experiment. Rooms were controlled for temperature (21 ± 1°C) and photoperiod (12:12 light/dark) for the entire study. All procedures were approved by the Iowa State University Institutional Animal Care and Use Committee and adhered to NIH guidelines.

### 2.2 Experimental Design

Upon arrival, twenty-four rats were individually housed in a standard laboratory cage with (Runner) or without (Sedentary) a locked 13-inch diameter running wheel (STARR Life Sciences, PA, USA) for one week. Note that rats in the sedentary group were deliberately not housed in cages with locked running wheels to keep physical activity to a minimum, as rodents tend to climb on locked wheels^47,48^. Half of the rats in each exercise condition were then randomly assigned to receive a single episode of 0 (no stress) or 100 (stress) uncontrollable tail shocks (**Fig. 1A**). The tail shock paradigm used in the present study followed our previous publications^24,49,50^. Approximately 2-5 h into the light cycle, rats assigned to the stress condition were restrained in flat bottom Plexiglas tubes with the tail protruding from the back where electrodes were placed to deliver 100, 1.25mA average current, 5-s tail shocks on a variable 1 min inter-shock interval. This procedure lasted approximately 100 minutes, though the precise time varied^24,49,50^. Rats that did not receive stress (i.e. 0 shocks) remained undisturbed in home cages in a different room during the tail shocks.

**Figure 1.**
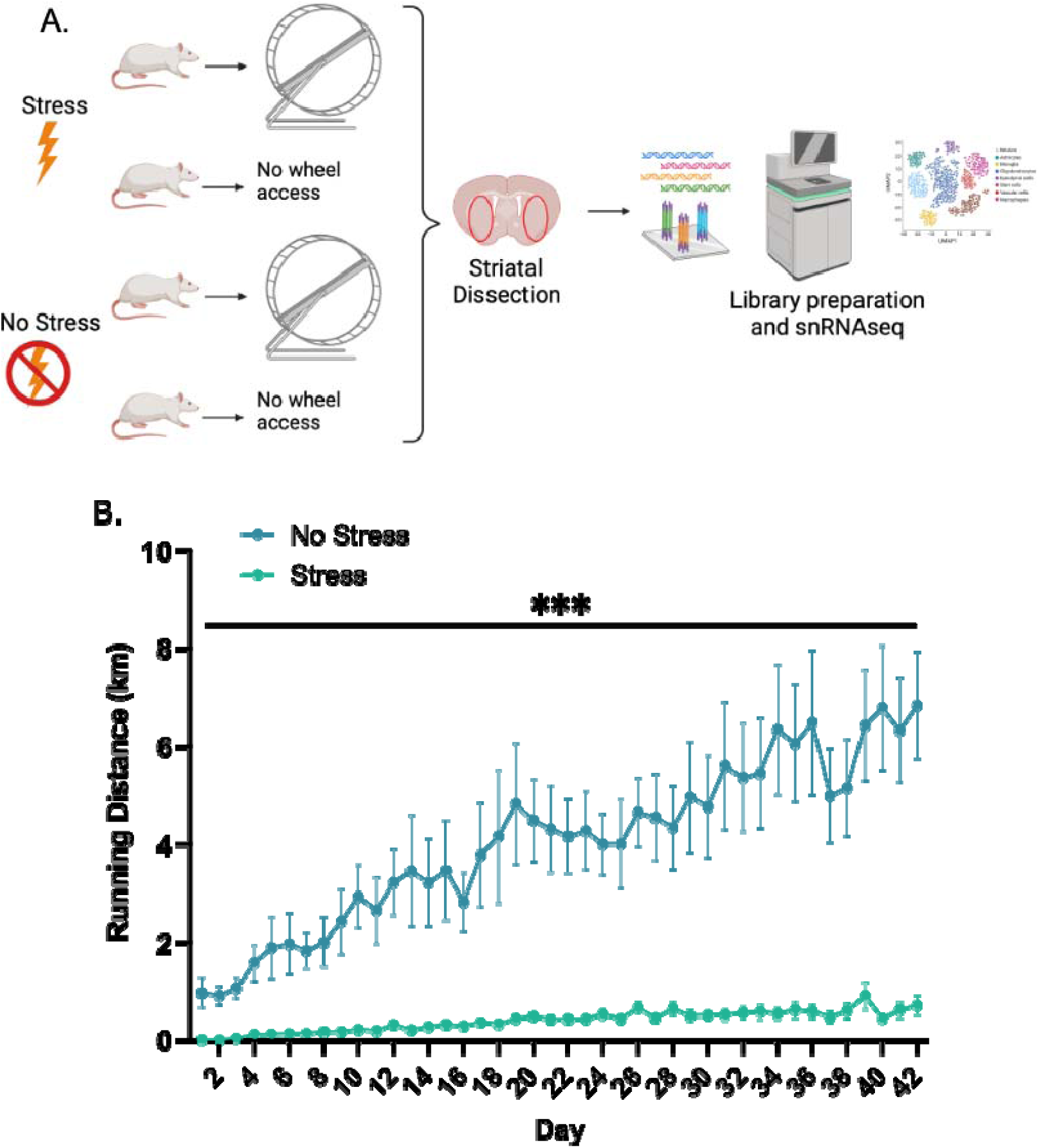
Persistent decreases in voluntary wheel running behavior are evident six weeks after an inescapable and acute stressor. (**A**) Male rats were split into four groups (n=6 per group) that received either one bout of inescapable stress or not and were then given access to a running wheel or not. Six weeks later, the striatum was dissected for single-nucleus RNA-sequencing (snRNAseq) (created with BioRender.com). (**B**) Stressed rats ran significantly less than non-stressed rats over the entire 6-week period (marked by ***; p<0.001).

48 hours later, running wheels of rats in the runner group were unlocked and rats were allowed to run freely for the six-week duration of the study. The 48-hour delay between tail shocks and wheel access was done to minimize risk of stressed rats developing an association between tail shock exposure and free access to running wheels. Wheel running distance was measured every hour for the entire six-week duration of the experiment. This time period was chosen because six, but not three-, weeks of running wheel access can prevent the development of striatum-involved anxiety– and depression-like behavior in rats^51–53^. Moreover, SD rats continue to display acute stress-induced persistent reductions in voluntary wheel running behavior beyond six weeks^11,12^. On Day 42, all subjects were euthanized by rapid decapitation during the peak running activity period (between 1-5 hours into the dark cycle). The final groups and sample sizes were sedentary no stress (n=6), sedentary stress (n=6), runner no stress (n=6), and runner stress (n=6).

In addition to measuring wheel revolutions continuously in 1-hour intervals in the rats that had access to running wheels throughout the entire experiment, we also measured body mass at the beginning and end of the experiment. We measured food consumption on days 21, 23, 25, 27, and 41 by subtracting the total weight of the food pellets in each cage from the prior day’s weight. These outcomes were measured to investigate whether stress caused long-term hypophagia.

### 2.3 Behavioral Analysis

The wheel running data (total revolutions over the entire 42-day experiment) were analyzed using a repeated-measures ANOVA with day as the within-subjects factor and treatment group (stress or no stress) as the between-subjects factor. The differences in body mass between the last day of the experiment and the first day were analyzed using a 2×2 ANOVA, with stress, exercise, and their interaction as factors. The food consumption data were analyzed using a 2×2 repeated-measures ANOVA with the 5 sampling days as the within-subjects factor. When interactions were significant, pairwise differences between means were evaluated using least-significant difference tests.

### 2.4 Nuclei Isolation, Library Construction, and Sequencing

Rat brains were rapidly extracted. The striatum was microdissected on a glass plate over ice per our previous publications^49,54,55^. The striatum was placed in pre-weighed cryovials, flash-frozen with liquid nitrogen, and then stored at −80°C until nuclei isolation. For nuclei isolation, the striata from all six rats within each treatment group (i.e. no stress sedentary, no stress runner, stress sedentary, or stress runner) were pooled together and placed in a centrifuge tube, lysed, and dissociated together to yield four samples. The nuclei were extracted and counted following protocols described previously^56^. Briefly, frozen striatal tissue was placed in lysis buffer (10 mM Tris-HCL, 10 mM NaCl, 3 mM MgCl_2,_ 0.1% Nonidet P40 Substitute) on ice for 30 min to lyse tissue followed by addition of Hibernate AB complete media (2% B27 and 0.05mM Glutamax, BrainBits HAB-500) and homogenization using a hand-held electronic homogenizer. Pooled homogenates were then centrifuged for 5 min at 600 x g and 4°C. The supernatant was discarded, and sample pellets were washed, resuspended, and then strained twice with 40uM strainers to remove any clumped cells or debris. The nuclei were then stained with 4’,6-diamidino-2-phenylindole (DAPI) and flowed through a fluorescence activated cell sorter (FACS) (BD FACSCanto, BD Biosciences, Franklin Lakes, NJ, USA) to assess viability. The nuclei were sorted according to their expression of DAPI and roughly 400,000 viable nuclei were collected per group. Following successful collection of the nuclei, the samples were given to the Iowa State DNA Facility who were responsible for making cDNA libraries using the 10x Chromium kit (3’ RNA library prep) (10X Genomics, Pleasanton, CA, USA), and subsequent sequencing on an S1 lane using the NovaSeq 6000 (Illumina, San Diego, CA, USA).

### 2.5 snRNA-seq Data Analysis

After sequencing, Raw FASTQ files were aligned to the *Rattus norvegicus* genome using the CellRanger pipeline (10X Genomics, Pleasanton, CA, USA). In total, 900,107,250 reads were sequenced from an estimated 44,629 nuclei. Across pools, the median number of Unique Molecular Identifiers (UMIs) sampled per nucleus ranged from 1,877-2,197, and the median number of genes sampled per nucleus ranged from 1,200-1,336. Filtered feature matrices generated by CellRanger were further pre-processed to 1) minimize risk of analyzing dead or dying nuclei by excluding barcodes associated with extremely few RNA molecules (<750) and genes (<500), 2) minimize risk of analyzing multiplets by excluding barcodes associated with an extremely large number of RNA molecules (>20,000) and genes (>5,000) suggesting multiple nuclei were sampled in the same droplet, 3) excluding barcodes associated with extremely high mitochondrial gene content (>5% of all genes). Individual rats were recovered from each pooled sample using the demultiplexing algorithm Souporcell^57^. Souporcell operates without the need for a reference genome for each individual by identifying putative individual-specific single-nucleotide polymorphisms (SNPs) associated with droplet-specific barcodes, and then using these SNPs to separate barcodes predicted to come from distinct individuals. Additionally, Souporcell uses these SNPs to estimate a probability that each droplet-associated barcode reflects multiple nuclei derived from genetically distinct individuals. As an additional quality control measure, we excluded those barcodes that were predicted by Souporcell to come from multiple individuals.

### 2.6 Clustering and Identification of Microglial Cluster

A data-driven clustering approach was used to dimensionally reduce the data and assign nuclei to communities based on transcriptomic similarity in R using Seurat ^58^. First, we normalized the data and identified variable genes using the ‘SCTransform’ function from the sctransform package in R. For dimensionality reduction, we performed Principal Components Analysis using the ‘RunPCA’ function (with dim=50) and Uniform Manifold Approximation (UMAP) using the ‘RunUMAP’ function (dims=1:50, min.dist=0.5, spread=0.2, n.neighbors=50, n.epochs=1000, metric=”euclidean”); both functions can be found in the Seurat package. Lastly, nearest neighbors were identified using the ‘FindNeighbors’ function (reduction=”pca”, k.param=50, dims=1:50, n.trees=500, prune.SNN=0) and clusters were assigned using the ‘FindClusters’ function (resolution=0.2, algorithm=2) for Seurat.

This pipeline resulted in 17 clusters. To aid in cluster identification, each parent cluster was assigned a number (0-16) which can be visualized in UMAP space (**Fig. 2A**). The microglia cluster was identified using genes specific to microglia (*P2ry12*, *Aif1*, *Itgam*, and *Cx3cr1*; see **Fig. 2B**). Re-clustering of microglia followed the same general workflow as above. Since the number of subclusters within the microglia cluster was not known *a priori*, this number of clusters was estimated using a non-parametric graph-based approach^59^. The approach involves constructing a k-minimum spanning tree (K-MST), which embeds nuclei into a similarity graph structure such that an edge exists between nuclei if they have similar gene expression information. Since different cluster resolutions lead to different clustering assignments, we estimated the number of clusters that achieves the greatest separation of cell clusters across different resolutions. To achieve this, we calculated the standardized within-cluster edge-count for a grid of resolution values. Cluster resolutions that result in well-separated communities of nuclei will lead to larger within-cluster edge-counts. It follows that the number of clusters can be determined using the resolution that maximizes the within-cluster edge-count. This graph-based approach circumvents the curse-of-dimensionality that is common in distance-based approaches to estimate the number of clusters, such as the gap statistic^60^.

**Figure 2.**
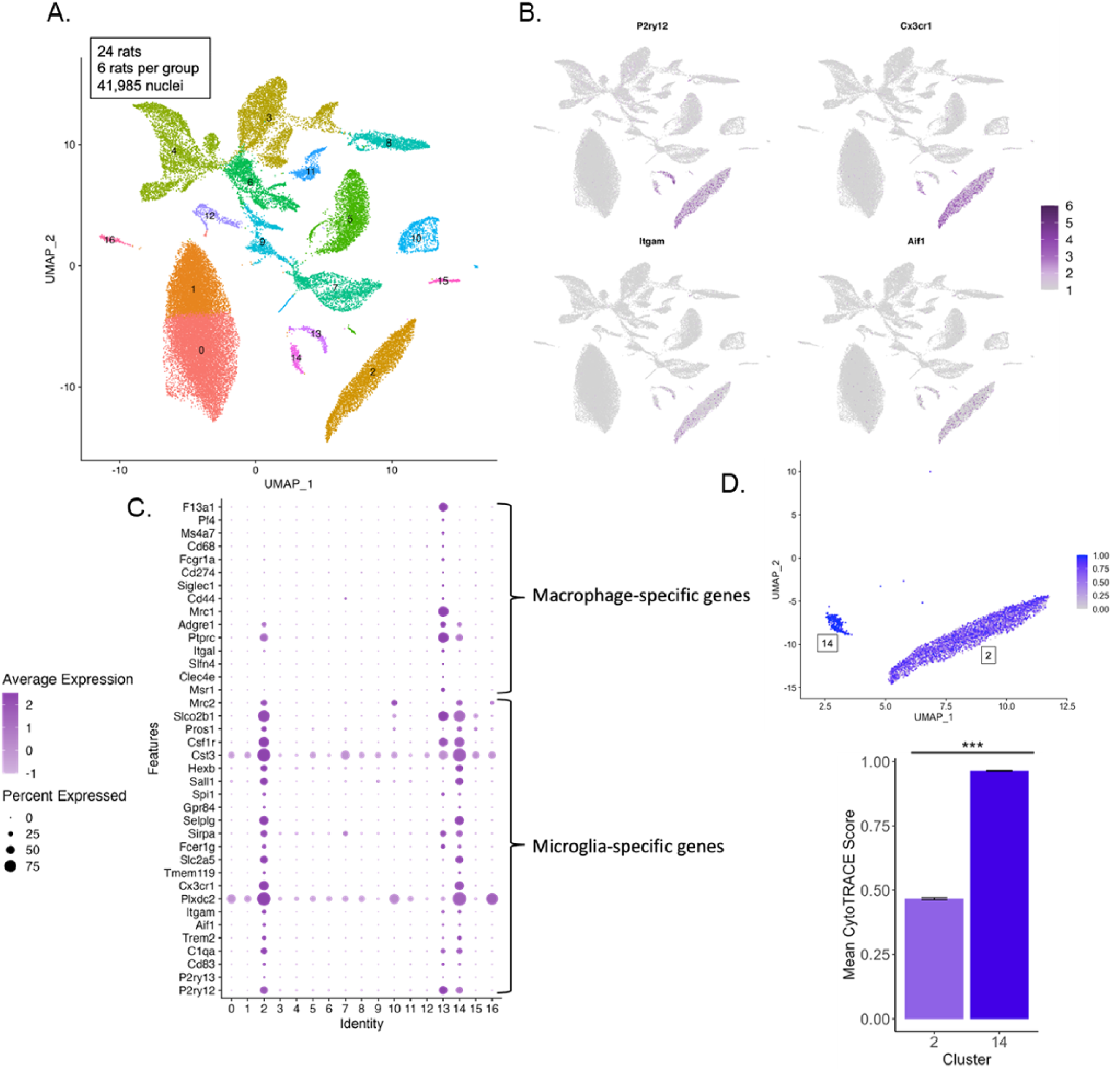
Identification and characterization of striatal microglia subtypes. (**A**) In total, 41,985 striatal nuclei were split into 17 distinct clusters based on their transcriptomes. (**B**) UMAP plots showing strong and selective expression of key microglia-related genes identified clusters 2, 13, and 14. (**C**) A dot plot showing divergent expression patterns microglia-specific and macrophage-specific genes in microglial clusters 2 and 14 versus macrophagic cluster 13. (**D**) CytoTRACE scores supported cluster 14 being composed almost exclusively of relatively immature microglia and cluster 2 containing a more diverse composition of intermediate maturity scores (marked by ***; p<0.001). The bars represent means of all the nuclei collapsed across all individuals, and standard errors reflect variation among nuclei not individuals.

### 2.7 Differential Gene Expression Analysis

For the microglia cluster, gene expression was compared between treatment groups and as a function of running distance using a negative binomial mixed effects model to accommodate for the multiple nuclei measured per individual. This was implemented using the ‘glmer’ package in R. The response variable was the number of transcripts of a gene per nucleus. Group (categorical variable) or running distance (continuous variable) was specified as a fixed effect and individual as a random effect. Dispersions were estimated using a Gamma-Poisson model within the ‘glmGamPoi’ R package^61^. The natural logarithm of the library sizes per nucleus was entered as an offset term to normalize variation in sequencing depth across the nuclei. To correct for multiple testing, we defined significance with an FDR cutoff of 0.05 using Storey’s q-value (q<0.05)^62^. To determine differentially expressed genes between specific subclusters, the Wilcoxon Rank-Sum approach, which requires no distributional assumptions, was used within Seurat’s ‘FindMarkers’ function. In addition to the above, we also analyzed gene expression a different way by taking the average gene expression using the SCtransform function (which corrects for library size) across all nuclei within individuals^63^. This metric was used for the mediation analyses, where we evaluated evidence for potential causal relations between stress, gene expression, and running levels using the ‘mediation’ R package^64^.

### 2.7 Gene Ontology

To test for overrepresentation of biological processes in differentially expressed genes (DEGs), we used the clusterProfiler package in R ^65,66^. GO terms were considered significant by Fisher’s Exact Test if FDR-corrected p<0.05. In the dot plots generated, GeneRatio is the ratio of input genes that are annotated in a term, count is the number of genes that belong to a given gene set, and the activated versus suppressed pathways are based on the normalized enrichment score (NES). NES is an indicator of the degree of enrichment. A positive NES indicates that members of the gene set tend to appear disproportionately at the top of a ranked input gene list, supporting pathway activation, and a negative NES indicates the opposite and is often considered an indication of pathway suppression ^65,66^.

### 2.8 Microglia Activation and Metabolism Scores

To determine changes in states of activation and function within our groups, a score was generated for the pro-inflammatory microglia state of activation (M1), the anti-inflammatory state of activation (M2), and microglial metabolism. First, we used sets of genes associated with M1, M2, and metabolic microglial states outlined in Ruan et al. (2023). The lists of these genes are in **Supplemental Table 12**. We then generated M1 and M2 scores by summing the expression (raw number of transcripts) of the predefined M1-associated and M2-associated genes, respectively, for each nucleus. Similarly, a metabolism score was calculated based on the sum of expression levels of key metabolic genes. These scores were normalized by the total transcript counts per nucleus to account for variations in sequencing depth. Statistical analyses, including linear mixed-effects models, were applied to examine the relationship between these scores, treatment groups, and microglia subclusters.

### 2.9 Cellular maturity

To estimate the relative age of microglial subclusters, CytoTRACE (Cellular (Cyto) Trajectory Reconstruction Analysis using gene Counts and Expression) was used^68^. CytoTRACE is an algorithm designed to predict the differentiation states of cells based on single-cell RNA sequencing data. The core principle of CytoTRACE is that less differentiated (immature) cells exhibit higher transcriptional diversity, whereas more differentiated (mature) cells show more specific and lower transcriptional activity. The algorithm operates through several key steps: it begins by normalizing the gene count matrix obtained from single-cell RNA sequencing data to ensure that variations in sequencing depth across cells do not bias the results. It then identifies genes with high variability across nuclei, as these are indicative of differentiation status. For each nucleus, CytoTRACE calculates a measure of transcriptional diversity, which involves computing the number of unique genes expressed and the expression levels of those genes^68^.

Based on this transcriptional diversity, CytoTRACE assigns a score between 0 and 1 to each cell, with a score of 1 indicating a cell that is predicted to be highly mature with broad gene expression patterns and a score of 0 indicating a cell that is predicted to be fully mature with more specialized gene expression patterns^68^. The CytoTRACE scores for the microglia were then analyzed using a linear mixed model, where the cluster (or subcluster) was treated as the fixed effect and individual was treated as the random effect. This statistical approach allows for the evaluation of cluster-specific effects while accounting for variability among individuals, facilitating estimation of differentiation states across clusters while controlling for potential confounding factors related to individual differences.

### 2.10 Nuclei Proportions

Differences in the relative proportions of microglial subclusters among groups were analyzed using a Chi-square test and Fisher’s Exact Test for pairwise comparisons. Significant group differences were defined using a significance threshold of alpha=0.05.

## 3 Results

### 3.1 Acute stress persistently impairs running behavior

The control rats that had access to running wheels (not stressed; **Fig. 1A**) displayed typical running patterns for Sprague Dawley males, including daily running distances of approximately 1-7 km that increased over the course of the experiment (42 days)^69–71^ (**Fig. 1B**). This increase over time was supported by analysis of running distances across days, which revealed a main effect of day on running distance (F(41, 410)=10.4, p<0.001). Analysis of total running distances between groups over the course of the experiment revealed a significant main effect of stress on running distance (F(1, 10)=24.4, p<0.001), a pattern that was driven by stressed rats running significantly shorter distances than rats that were not stressed (Fig. 1B). Notably, the difference in running distance between stressed and non-stressed animals increased across the days (stress x day interaction, (F(41, 410)=6.5, p<0.0001). When collapsing running distances across days, stressed animals ran on average 14.9% as far as non-stressed animals (**Fig. 1B**). Results for body mass and food intake for all rats are reported in Supplementary Results and shown in **Supplementary Figure 1A-B**.

### 3.2 Identification of microglia clusters

The clustering algorithm identified 17 clusters based on single nucleus transcriptomes (**Fig. 2A**). To determine if specific clusters contained nuclei derived from microglia, we examined expression patterns of several canonical microglial marker genes (*P2ry12*, *Aif1*, *Itgam*, and *Cx3cr1*) across clusters (**Fig. 2B**). All four genes showed strong and selective expression in just three clusters (clusters 2, 13, and 14). Analysis of marker genes in each cluster using the Seurat ‘FindAllMarkers’ (using a Wilcoxon Rank Sum test) function showed strong expression of canonical markers of microglia and macrophages within these three clusters as well.

To more closely examine these clusters, we curated lists of microglia-specific (*P2ry12, P2ry13, Cd83, C1qa, Trem2, Aif1, Itgam, Plxdc2, Cx3cr1, Tmem119, Slc2a5, Fcer1g, Sirpa, Selplg, Gpr84, Spi1, Sall1, Hexb, Cst3, Csf1r, Pros1, Slco2b1, Mrc2*) and macrophage-specific marker genes (*Msr1, Clec4e, Slfn4, Itgal, Ptprc, Adgre1, Mrc1, Cd44, Siglec1, Cd274, Fcgr1a, Cd68, Ms4a7, Pf4, F13a1*) (**Supp. Fig. 2**). Clusters 2 and 14 showed strong and selective expression of microglia-specific marker genes (**Supp. Fig. 2A**), whereas cluster 13 showed strong and selective expression of macrophage-specific marker genes (**Supp. Fig. 2B**). Additionally, we plotted the same set of microglia-specific genes as well as macrophage-specific genes. This analysis again revealed a clear pattern whereby cluster 13 showed strong and selective expression of macrophage-specific genes while clusters 2 and 14 showed strong and selective expression of microglia-specific genes (**Fig. 2C**).

Based on these gene expression patterns and presence of these marker genes within each cluster, we therefore considered two clusters (clusters 2 and 14) as containing nuclei derived from microglia. To better understand the biological differences between these two microglial clusters transcriptomic signatures of cell maturity were investigated using CytoTRACE, a bioinformatics tool that assigns a differentiation score to each nucleus based on transcriptional diversity, with a greater score indicating a less differentiated nucleus (see Methods section 2.8). One cluster (cluster 14) showed a significantly greater CytoTRACE score compared to the other (cluster 2; β = .47, SE= .013, p<.0001), supporting the idea that it consisted of relatively less differentiated microglia while the other cluster (cluster 2) consisted of relatively more mature microglia (**Fig. 2D**) (**Supp. Table 1**). Thus, for the remainder of this paper we focused our analysis on the relatively mature microglia from cluster 2.

### 3.3 Striatal microglial signatures of stress

To investigate signatures of stress in striatal microglia, gene expression was compared between the non-stressed sedentary group (no stress control) and the stressed sedentary group six weeks after acute stress when running deficits are still present (**Fig. 1B**). In microglial cluster 2, 24 stress-related differentially expressed genes (stress-DEGs) were identified, of which 3 were upregulated in the non-stressed animals while the remaining 21 were upregulated in the stressed animals (**Fig. 3A**) (**Supp. Table 2**). A chi-square goodness-of-fit test revealed that the relative proportions of upregulated versus downregulated stress-DEGs in this cluster to be significantly different from 50:50, i.e., significantly more genes were upregulated versus downregulated in association with stress 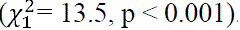. Several stress-DEGs that were upregulated in association with stress have previously been shown to reflect microglia activation, including *Cst3, Apoe, C1qa,* and *C1qb* ^72–77^ (**Fig. 3B**).

**Figure 3.**
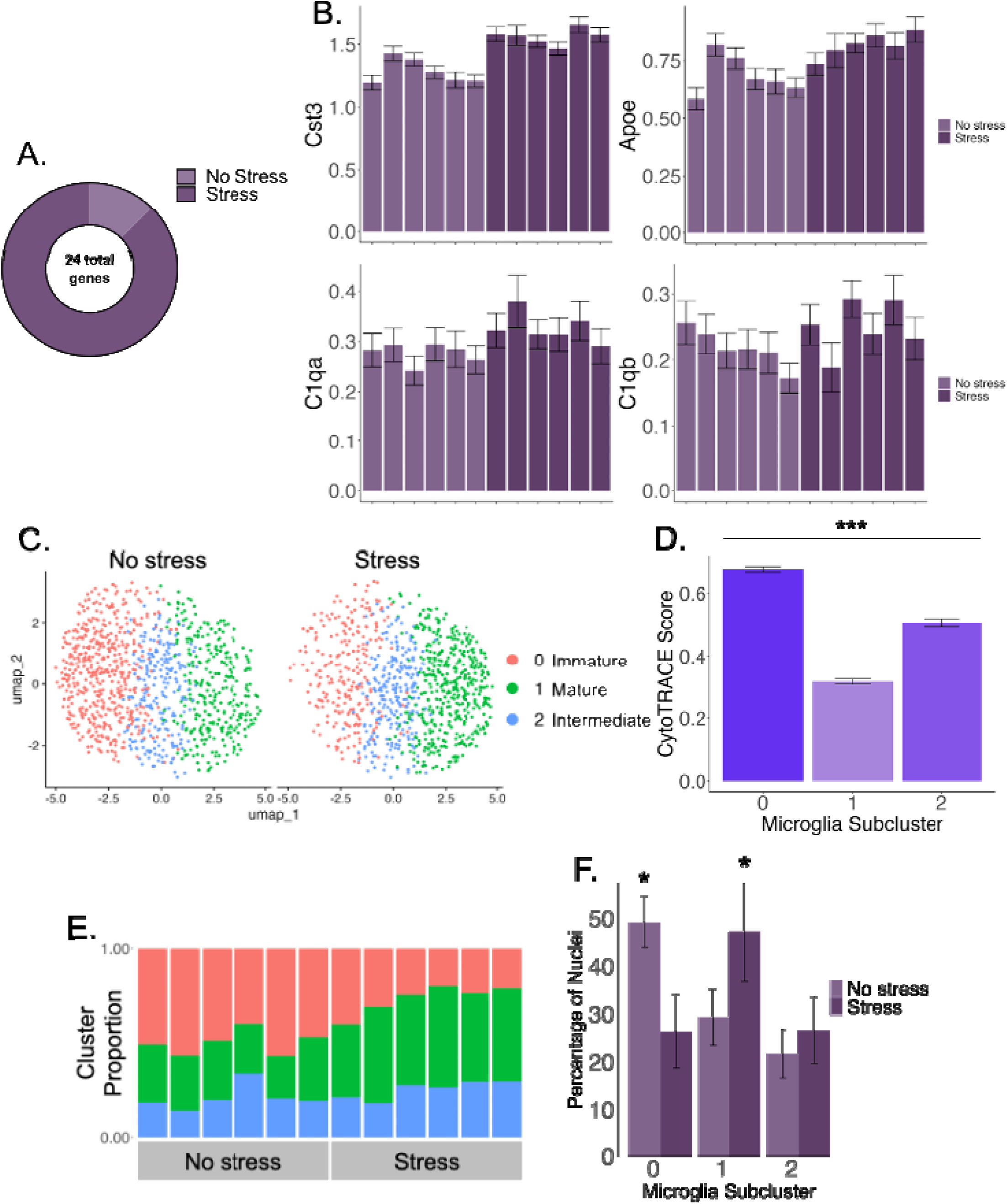
Acute stress produces prolonged striatal microglial activation. (**A**) A comparison of gene expression between the no stress and stress groups revealed 24 stress-DEGs. The majority stress-DEGs were upregulated in the stress group compared to the no stress group. (**B**) Mean gene expression (SCTransformed) of stress-DEGs related to microglial activation are shown. Individual means and standard errors (among nuclei) are shown for all individuals in the no stress and stress groups. (**C**) Re-clustering of microglia revealed three subclusters: immature (pink), mature (green), and intermediate (blue). (**D**) These subclusters differed in predicted levels of cellular maturity estimated by CytoTRACE. (**E**) The relative proportions of each subcluster are shown for each individual. (**F**) Individuals in the stress group displayed lesser proportions of relatively immature microglia and greater proportions of relatively mature microglia compared to individuals in the no stress group (marked with *; p<0.05).

To further evaluate differences between the non-stressed and stressed rat, parent cluster 2 (i.e., relatively mature microglia cluster) was re-clustered and showed three distinct sub-clusters (**Fig. 3C**). CytoTRACE analysis showed that re-clustering differed strongly in predicted cellular maturity. Based on CytoTRACE scores, subcluster 0 was the least differentiated, followed by subcluster 2 and then subcluster 1 (**Fig. 3D**) (**Supp. Table 3**). Thus, for the sake of clarity, we hereafter refer to subclusters 0, 2, and 1 as putatively immature, intermediate, and mature, respectively.

Significant stress-DEGs were also found when analyzing each of the 3 subclusters of microglia separately. Despite the reduced sample size of approximately 1/3 the number of nuclei per subcluster, 1 stress-DEG was detected in the immature cluster (**Supp. Table 4**), 4 in the mature subcluster (**Supp. Table 5**), and 4 in the intermediate subcluster (**Supp. Table 6**). All these genes were contained within the 24 stress-DEGs when considering all subclusters together. Overall, these findings suggest that stress changes gene expression in all subtypes of striatal microglia, though the majority of the effect is in the mature and intermediate subclusters.

The influence of stress on microglia maturation phenotypes was investigated next by comparing relative proportions of putatively immature, intermediate, and mature microglia differed between no stressed and stressed rats. Stressed rats displayed greater relative proportions of mature microglia and lesser relative proportions of immature microglia compared to the no stress rats (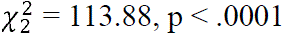; pairwise differences indicated stress differed from no stress in the relative proportions of immature and mature subclusters, p<0.05, but not in the relative proportions of the intermediate subcluster; **Fig. 3E & 3F**).

To investigate the possible relationship between stress, microglia maturity, and activation patterns, the expression of genes involved in glial activation and metabolism was scored for each nucleus. M1 is considered a canonical pro-inflammatory pattern of activation that microglia can adopt while M2 is a canonical anti-inflammatory pattern adopted by microglia^78^. Furthermore, studies have shown that microglia polarization to an M1 phenotype is linked to metabolic shifts from oxidative phosphorylation to aerobic glycolysis, influencing their functional activity thereby highlighting the importance of generating a metabolism score^78^. However, the M1, M2, and metabolism scores did not significantly differ among the putatively immature, intermediate, and mature microglial subclusters. Subsequent analysis further revealed that none of these scores differed between the stress and no stress rats (**Supp. Fig. 3A-C**). Taken together, these findings suggest that canonical M1/M2 signatures do not account for the major differences observed between groups when the data are restricted to non-runners.

### 3.4 Striatal microglial signatures of exercise enrichment

We next investigated the impact of exercise enrichment on striatal microglia gene expression. Exercise enrichment, defined as access to a running wheel (independent of how much the animal runs), was first studied in the absence of stress (i.e., this analysis was done only on the no stress sedentary and runner groups). For the purposes of this analysis, we define the no stress runner group as the ‘exercise enrichment’ group. We found 201 significant differentially expressed genes between the no stress sedentary and exercise enrichment groups (exercise enrichment-DEGs). A disproportionate number of exercise enrichment-DEGs were upregulated in the sedentary group (n=158) versus the exercise enrichment group (n=43; chi-square goodness-of-fit test, 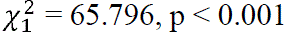) (**Fig. 4A**) (**Supp. Table 18**). Examples of exercise-enrichment DEGs that were upregulated in the sedentary group and exercise-enrichment DEGs that were upregulated in the exercise enrichment group are shown in **Figure 4B**. Specific GO terms that were significantly over-represented among exercise enrichment-DEGs included: cellular respiration/oxidative phosphorylation (e.g., *Atp7a, Cox5a*), immune response (e.g., *Cd74, Tnf*), synaptic signaling (e.g., *Camk2a, Sorcs3*), and neurogenesis (e.g., *Fgf13, Nrp1*) (**Fig. 4C**) (**Supp. Table 20**). These results are consistent with exercise enrichment shifting striatal microglia toward functions related to synaptic signaling and remodeling via phagocytosis of neuronal processes.

**Figure 4.**
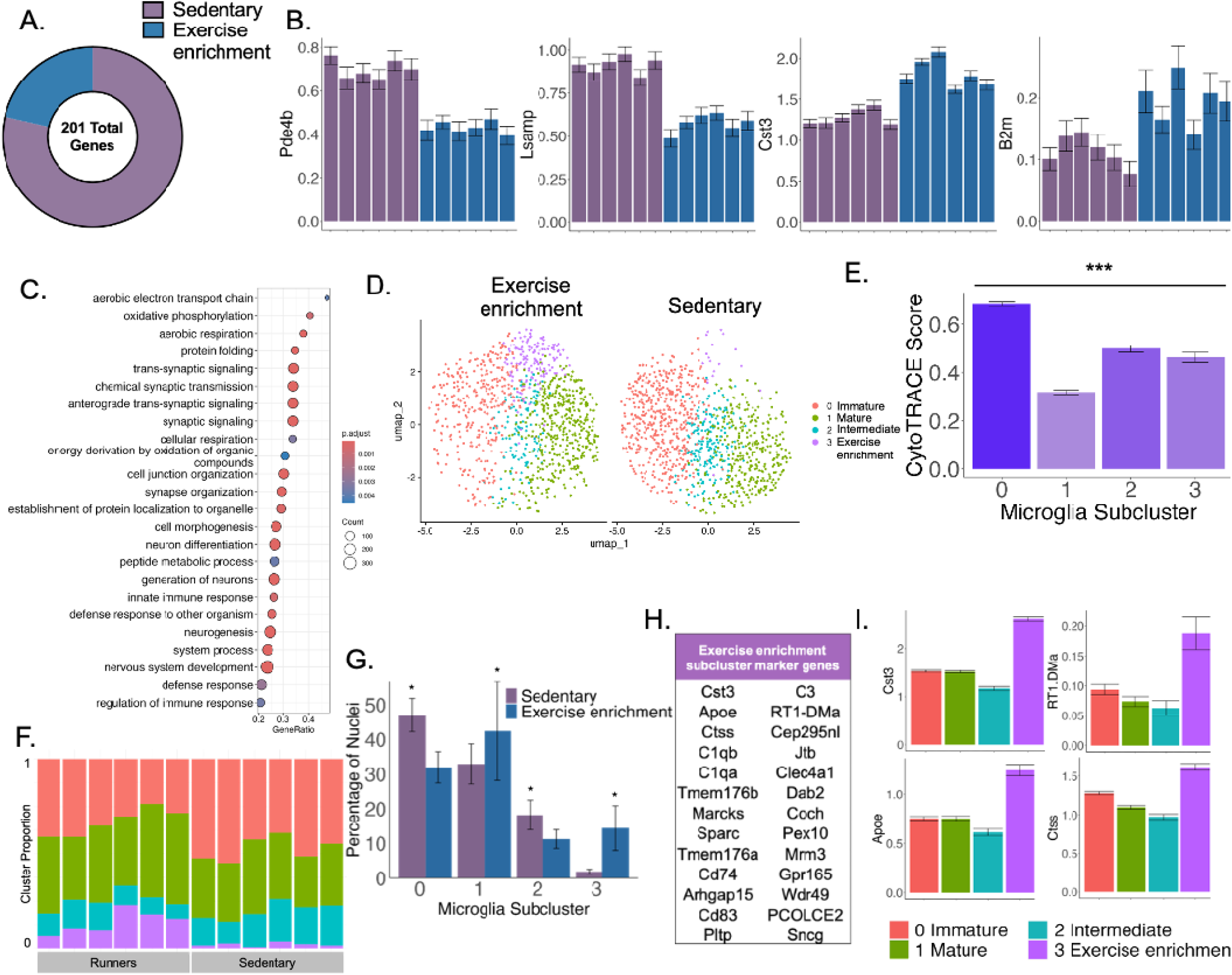
Exercise enrichment is associated with a specific striatal microglial activation profile. (**A**) A comparison of gene expression between the sedentary and exercise-enrichment groups found a total of 201 differentially expressed genes, or exercise enrichment-DEGs, the vast majority of which were upregulated in the sedentary group relative to the exercise enrichment group (**B**) Mean gene expression of exemplar genes upregulated in the sedentary and exercise enrichment groups are shown. Individual means and standard errors (among nuclei) are shown for all individuals in the sedentary and exercise enrichment groups. (**C**) Dot plot highlighting GO terms that were significantly enriched among exercise enrichment-DEGs. (**D**) Re-clustering identified four distinct subclusters of microglia in the sedentary and exercise enrichment groups, including a fourth subcluster that was abundant in exercise enriched but not sedentary rats. (**E**) CytoTRACE analysis revealed a similar maturity structure as the stress vs. no stress analysis (subcluster 0 = immature, subcluster 1 = mature, and subcluster 2 = intermediate), with the additional exercise enrichment subcluster showing a relative maturity level similar to the intermediate subcluster. (**F**) Proportions of nuclei assigned to each subcluster are shown for each of the individuals. For legend see panel I. (**G**) Individuals in the exercise enrichment group displayed a significantly lower proportion of relatively immature microglia, a greater proportion of relatively mature microglia, and an even greater proportion of microglia assigned to the exercise enrichment subcluster, which was nearly absent in sedentary rats (marked with *, p<0.05). (**H**) Comparison of the exercise enrichment subcluster to the other subclusters revealed a distinct set of genes related to microglial activation, phagocytosis, and the complement system. (**I**) Expression patterns of example genes that were upregulated in the exercise enrichment subcluster.

To further analyze the effects of exercise enrichment on microglia, parent cluster 2 was re-clustered containing only the exercise enrichment and sedentary no stress groups. This analysis revealed the same three subclusters derived in Fig. 3C, along with a fourth distinct subcluster (**Fig. 4D**). The majority of the cells in this fourth subcluster were from the exercise enrichment group (**Fig. 4D**) (**Supp. Fig. 4**). Indeed, within the exercise enrichment group, 14.4% of striatal microglia (in parent cluster 2) were assigned to the exercise enrichment subcluster, whereas only 1.9% of striatal microglia were assigned to this subcluster in sedentary rats. CytoTRACE suggested that the remaining three subclusters were distinguished by relative maturity, mirroring the pattern observed in Fig 3D. Thus, these subclusters are again referred to as the immature, intermediate, and mature subclusters. The fourth subcluster is referred to as the ‘exercise enrichment’ subcluster hereafter for the sake of clarity. Notably, the exercise enrichment subcluster and the intermediate subcluster displayed similar CytoTRACE scores, supporting that they were both composed of relatively intermediate microglia (**Fig. 4E**) (**Supp. Table 8**).

The relative proportions of the four microglial subclusters differed between exercise enriched and sedentary groups 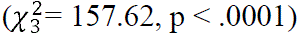 (**Fig. 4F & 4G**). The greatest difference was for the exercise enrichment subcluster, as described above, where the proportion of this subtype of microglia was 7.6-fold higher in runners than the sedentary group. The exercise enrichment group also had significantly fewer intermediate and immature microglia, and significantly more mature microglia compared to the sedentary rats (Fig. 4F & 4G) (**Fig. 4F & 4G**). Differential gene expression and gene set enrichment analyses further revealed that the genes upregulated in the exercise enrichment subcluster relative to the other subclusters were generally related to microglia activation/immune response (*Cst3, Marcks, Sparc, Cd83, Dab2, Tmem176a, Tmem176b, Clec4a1, and Cd74*), the complement system (*C1qa, C1qb, C3*), and microglia phagocytosis (*Ctss, Pltp, RT1-DMa*) (**Fig. 4H**). Lastly, differences among subclusters were analyzed for M1, M2, and metabolism scores described above (**Supp. Table 12**)^67^. Similar to our earlier analyses, no significant differences between exercise enrichment and sedentary groups or among subclusters were observed (**Supp. Fig. 3D-F**). Taken together, these results suggest that canonical signatures of microglia activation do not sufficiently describe subtypes of striatal microglia or their responses to exercise enrichment in our paradigm.

### 3.5 Striatal microglial signatures of individual running levels

To further investigate the regulation of gene changes by wheel running, we analyzed gene expression associated with voluntary wheel running distances in non-stressed rats across multiple timescales relative to when the striatum was sampled: the previous 40 days, the previous day, the previous three hours, the period 30-90 minutes before sampling, and the 30-minute period immediately before sampling (**Supp. Table 9**; **Supp. Tables 13-16, respectively**). Strikingly, 807 genes showed expression associated with total distance run over the entire 6-week period (6-week distance-DEGs; q<0.05), but far fewer distance-DEGs were observed when analyzing total running distances over the final 24 hours (n=58 distance-DEGs), 3 hours (n=20 distance-DEGs), 30-90 minutes (n=20 distance-DEGs), and 30 minutes (n=9 distance-DEGs). Notably, the most of the distance-DEGs identified in analyses of these shorter time intervals were also identified in our analyses of the entire 6-week period, likely due to the positive correlations between 6-week running distances 6-weekand running distances over the smaller intervals (Pearson’s R was 0.47, 0.14, 0.1, and 0.03 for the correlation between running distances over 40 days versus running distances over 24 hours, 3 hours, 30-90 min, and 30 min respectively). Interestingly, there was little overlap between running level-DEGs and exercise enrichment-DEGs as indicated by hypergeometric test (p=0.66), suggesting that different genes are regulated by running levels versus running wheel access in striatal microglia (**Supp. Fig. 5A, example relationships between distance-DEG expression and running distance shown in Supp. Fig. 5B**). Taken together, these results support that striatal microglial gene expression most strongly reflects long-term as opposed to recent levels of running.

### 3.6 Striatal microglial signatures of interactions between stress and exercise

To investigate possible interactions of acute stress and exercise on striatal microglial gene expression, we next analyzed all four experimental groups using a 2×2 design with stress and exercise as main effects as well as a stress by exercise interaction term. This analysis revealed 207 exercise-DEGs (main effect of exercise), 201 stress-DEGs (main effect of stress), and 17 interaction-DEGs (**Supp. Table 10, example interaction-DEGs are shown in Fig. 5A**). Stress-DEGs and exercise enrichment-DEGs in this analysis overlapped significantly with the stress-DEGs and exercise enrichment-DEGs identified earlier when stress and exercise were analyzed separately (**Supp. Fig. 6A & 6B**; hypergeometric tests, both p<0.05). There was no significant overlap between stress-DEGs and exercise-DEGs from either analysis (hypergeometric test on stress-DEGs and exercise-DEGs from analyses using a 2×2 model and analyses of each factor separately (**Fig. 5B & 5C**). Taken together, these results suggest that stress and exercise have long-term and robust effects on expression of largely distinct genes in striatal microglia.

**Figure 5.**
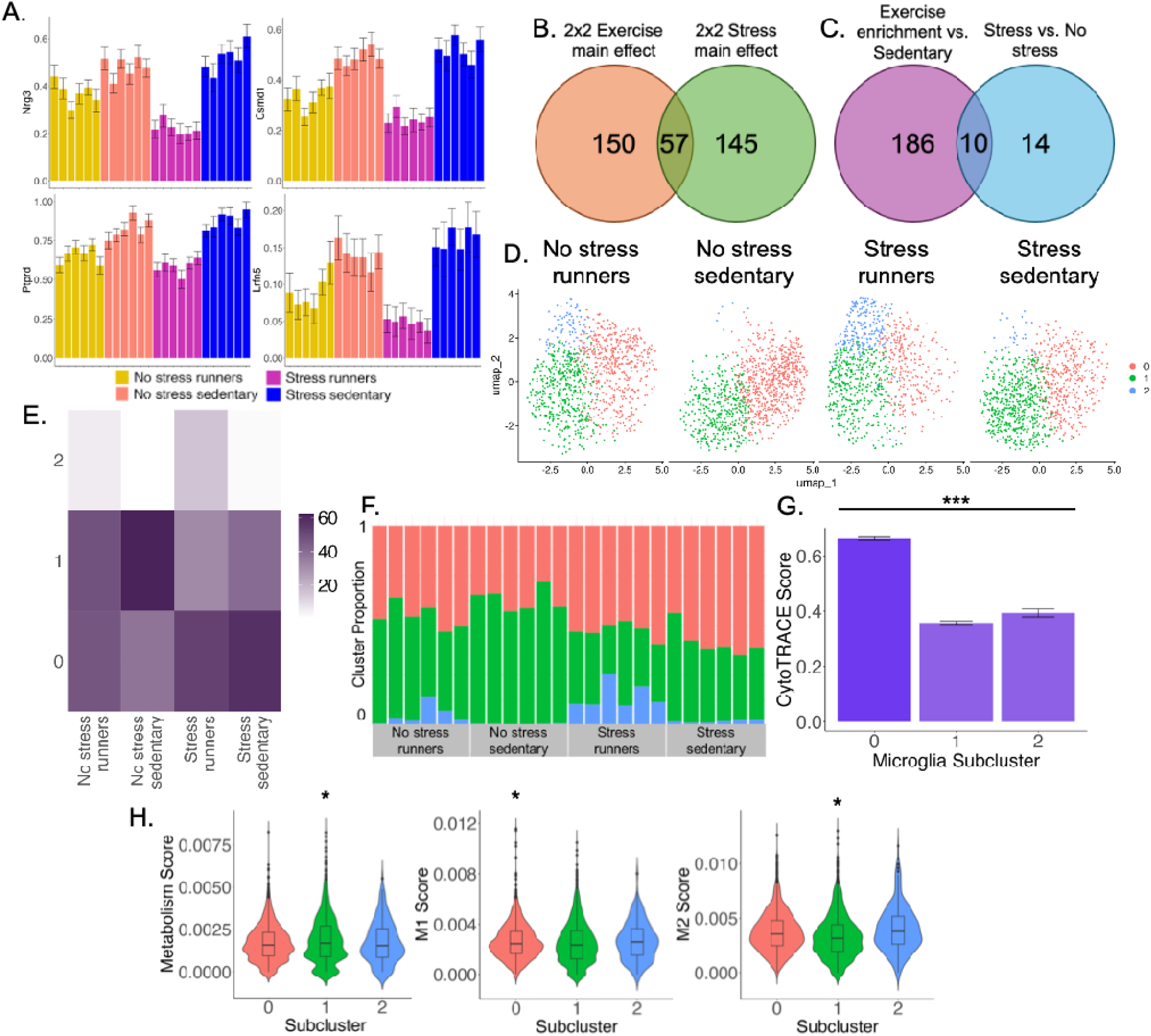
Striatal microglial signatures of stress-by-exercise interactions. (**A**) Analysis of all four groups yielded 207 significant exercise-DEGs (main effect of exercise), 201 stress-DEGs (main effect of stress), and 17 interaction-DEGs. Mean normalized gene expression for four exemplar interaction-DEGs are plotted. Individual means and standard errors (among nuclei) are shown. (**B**) Stress-DEGs and exercise enrichment-DEGs did not significantly overlap (results shown reflect main effects of stress and exercise in the 2×2 analysis). (**C**) Similarly, stress-DEGs and exercise enrichment-DEGs from the separate analyses did not significantly overlap. (**D**) Re-clustering of microglia from all treatment groups resulted in three distinct subclusters, including a subcluster that is only apparent in runners (subcluster 2). (**E**) Proportions of nuclei in each subcluster are shown for each treatment group. (**F**) The runner subcluster accounted for a large percentage of microglia in all individuals that had access to running wheels (both stress and no stress), regardless of running levels, but was almost entirely absent in individuals without running wheels. (**G**) Subcluster 1 was relatively more immature than the other clusters (marked by ***; p<0.001). (**H**) The subclusters were differentiated based on genes related to metabolism, pro-(M1), or anti-inflammatory (M2) activation. Specifically, the runner subcluster showed a lower metabolism score and higher M2 score compared to the relatively mature subcluster 1.

We additionally ran an analysis of covariance (ANCOVA) to investigate how stress and long-term running distance may interact to influence striatal microglial gene expression determine (**Supp. Table 17**). This analysis included only animals housed with a running wheel, both stressed and not stressed. Results revealed 401 6-week distance-DEGs, 379 stress-DEGs, and 355 significant stress-by-distance interaction-DEGs. Distance-DEGs identified in the ANCOVA overlapped significantly with distance-DEGs identified in earlier analyses of the non-stressed runners only (hypergeometric test, p < 0.001; **Supp. Fig. 5D**). Interestingly, stress-DEGs identified in earlier analyses of stress versus no-stress (all sedentary) (**Supp. Table 2**) did not significantly overlap with stress-DEGs identified in the ANCOVA (p>0.05) (**Supp. Fig. 6C**), a pattern that was likely driven by differences in the effects of stress on striatal microgial gene expression between sedentary and exercising rats. Examples of genes that showed strong effects of the running covariate, a strong effect of stress, and no interaction are shown in **Supplementary Figure 6E & 6F**. Examples of genes that showed a strong interaction between the running covariate and stress are shown in **Supplementary Figure 6G**. Taken together, these results support an extraordinary degree of sensitivity to complex interactions between stress and exercise in striatal microglia.

Genes that showed a significant effect of the running covariate, main effect of stress or interaction in the ANCOVA analyses described above were further analyzed by mediation analysis. The purpose was to identify candidate genes involved in causing the reduced running behavior and other genes that are more likely a consequence of the different running levels rather than a cause. This analysis found 12 genes whose expression levels were significant mediators of the effects of stress on running levels, i.e., putative genes that cause varying running levels. The analysis also found 16 genes whose expression levels were mediated by different running levels, i.e., putative genes that change in response to stress as a consequence of reduced running (**Supp. Table 21**) (**Supp. Fig. 9**).

Lastly, we extended our analyses of microglial subclusters by re-clustering parent cluster 2 across all four groups. This analysis yielded three distinct subclusters (**Fig. 5D**). Mirroring our analyses of running rats only, one subcluster (subcluster 2) consisted of nuclei derived predominantly from runners (**Fig. 5E & Fig. 5F**). Analysis of cell maturity using CytoTRACE supported that subcluster 1 was relatively immature whereas subclusters 0 and 2 (the exercise enrichment subcluster) were relatively and comparably mature (**Fig. 5G**) (**Supp. Table 11**). The proportion of nuclei in each subcluster differed strongly among treatment groups in an overall Chi-square test (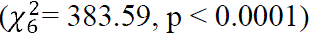). Within each subcluster, all pairwise comparisons between treatment groups were significant as indicated by Fisher Exact Test (**Fig. 5E**). To investigate whether these subclusters differed in additional biological dimensions, we analyzed the same M1, M2, and metabolism scores described above (**Supp. Table 12**). This analysis revealed differences among subclusters: compared to the immature subcluster 1, the mature subcluster 0 displayed greater metabolism scores and decreased expression of both M1 and M2 genes, whereas the exercise-enrichment subcluster 2 displayed decreased expression of M1 genes (**Fig. 5H**). Within subclusters, no significant differences between treatment groups were observed for the M1, M2, or metabolism scores. Taken together, these results further reinforce the conclusion that the canonical M1/M2/metabolism metrics are not sufficient to differentiate the treatment groups, though with the full data set, the subclusters can be differentiated. The fact that the exercise-enrichment subcluster displays a higher M2 score than the relatively mature subcluster is indicative of a protective microglial functional state.

## 4 Discussion

The aim of this study was to define the molecular signatures of stress, exercise, and their interaction in striatal microglia. Using snRNA-seq, we showed stress-associated differences in estimated microglial maturity 42 days post-stressor as well as molecular signatures of a distinctive and non-canonical activation pattern, together underscoring the enduring impacts of stress on microglial dynamics. Access to a running wheel was also associated with profound alterations in striatal microglia, including greater relative proportions of a distinct microglial subpopulation that was nearly absent in sedentary rats. In this subpopulation, exercise enrichment was associated with increased expression of genes related to complement system– and phagocytosis-related pathways, indicative of a separate, distinct form of microglial activation. Even more striking, we observed distinct microglial gene expression signatures associated with access to a running wheel (i.e., exercise enrichment) compared to individual running distances. Lastly, we observed evidence for a distinct form of activation in response to both stress and exercise, potentially reflecting molecular processes that contribute to deficits in motivation for physical activity. Our findings thus support a broader spectrum of microglial functional states than previously appreciated and reveal candidate molecular pathways by which striatal microglia may regulate motivation for exercise following stressors.

We observed that a single session of inescapable foot shock was associated with severely depressed levels of voluntary wheel running behavior for 42 days after the stressor (**Fig. 1B**). This extends previous literature where reduced voluntary running was observed up to 28 days after the stressor^11,12^. We hypothesized that microglia-induced neuroinflammation and/or priming of microglia in the striatum may contribute to the reduced motivation for running given the role of microglia in both stress and exercise (as reviewed in ^38,79^). Consistent with our hypothesis, several genes that have been implicated in microglia activation were upregulated in the stress group: *Cst3, Apoe, Srsf7, Rpl19, C1qa,* and *C1qb* (some of which are plotted in **Fig. 3B**). *Cst3* and *Apoe* were documented previously as markers of activated microglia^72^, disease-associated microglia^72,73^, or injury-associated microglia^74,75^. In contrast, *Srsf7* and *Rpl19* have been found to be downregulated in microglia in Alzheimer’s disease^80^ and are associated with homeostatic microglia^81^. Both *C1qa* and *C1qb* are involved in the complement system which are indicators of microglia activation as well ^76,77^.

In addition to displaying differential expression of specific markers of microglia activation as described above, stress was also associated with greater estimated microglial maturity (**Fig. 3D & 3E**), a pattern that may reflect a specific form of inflammatory action. Ruan et al. (2023) used CytoTRACE in their experiment examining the role of leucine-rich alpha-2-glycoprotein 1 (Lrg1) in cerebral ischemia reperfusion injury and similarly found that different clusters of microglia had different states of maturation which they attributed to their inflammatory action. On the other hand, we did not see evidence of canonical markers of stress-induced inflammation such as *Il1β*, *Il6*, or *Tnfα* even though transcripts for these genes were detected in our dataset. Similarly, while we were able to detect expression of many genes that are considered markers of M1, M2, and metabolism states of microglia, these genes were not informative for distinguishing subclusters or effects of stress in sedentary rats (**Supp. Fig. 1A-F**). Together, this supports a unique form of enduring microglial activation following acute stress that is not captured by classic definitions of microglia activation (i.e., M1 and M2). This is consistent with previous work showing enduring responses of hippocampal and cortical microglia to inescapable shock stress^31,32,35,82^. Given the dual role of inescapable shock stress in producing both microglial activation and reduced motivation for exercise, it is possible that the reduced running we see in the present study is a result of the sustained striatal microglial activation following inescapable shock stress. Results of the mediation analysis support this idea and identify several candidate genes in microglia that may be partially responsible for regulating running behavior in response to stress (**Supp. Table 21**) (**Supp. Fig. 9**). Future studies are needed to determine if blocking striatal microglia activation ameliorates deficits in motivation for physical activity following exposure to a severe, acute stressor.

In addition to exhibiting molecular signatures of stress, microglia also exhibited molecular signatures of physical activity. Individuals with running wheel access that were not stressed ran every day throughout the experiment, and total running distance was associated with expression levels of 807 genes (**Supp. Table 9**). These effects were unlikely driven by acute effects of running, because more than 10-fold fewer running-level DEGs were observed when analyzing shorter intervals immediately prior to sampling compared to the total distance run over previous 40 days. This supports the idea that the cumulative effect of running over multiple days or weeks results in a robust functional shift in a subset of microglia, perhaps to support striatal function during running behavior.

In addition to running-level DEGs, we also detected a separate set of non-overlapping genes associated with running wheel access (**Supp. Fig. 4A**), supporting distinct microglial profiles associated with running wheel access versus levels of running. Access to a running wheel is considered a form of enrichment ^83–85^. Thus, we believe the DEGs identified in the comparison of runners to sedentary rats (no stress) reflect the enrichment of the running wheel, or the deprivation of no running wheel in the sedentary group, rather than differential levels of physical activity *per se*. Strikingly, we observed an ‘exercise enrichment’ subcluster that is only found in rats with access to a running wheel, whether stressed (and thus, running very low levels) or not (**Fig. 5D-F**). The genes distinguishing this subcluster from other microglial subclusters were generally related to microglial activation/immune response (*Cst3, Marcks, Sparc, Cd83, Dab2, Tmem176a, Tmem176b, Clec4a1, and Cd74*) the complement system (*C1qa, C1qb, C3*), and microglial phagocytosis (*Ctss, Pltp, RT1-DMa*) (**Fig. 4D**). Together, these results are consistent with microglia being activated and phagocytosing in a complement-dependent manner more in rats with voluntary wheel running enrichment compared to those deprived of the enrichment.

Activated microglia can eliminate both damaged cells and dysfunctional synapses, a phenomenon known as “synaptic stripping” ^76,86–89^. C1q and C3, integral components of the classical complement cascade, attach to the surface of invading pathogens. Subsequently, phagocytes expressing target complement receptors in the peripheral immune system eliminate these pathogens ^90,91^. Similarly, within the CNS, microglia utilize classical complement cascades to facilitate phagocytic signaling, leading to the removal of excess synapses ^76,77^. Few studies have evaluated microglial complement and phagocytosis during physical exercise, but persistent aerobic exercise has the potential to preserve the integrity and functionality of neurovascular units in aging mice^92^. Exercise achieves this by diminishing C1q-expressing microglia and enhancing neuroplasticity^92^. Moreover, physical exercise appears to facilitate cellular communication between microglia and neurons through complement molecules^93^. Thus, the emergence of a unique C1q+ and C3+ microglial subpopulation with exercise may be involved in neuron-microglia and/or astrocyte-microglia crosstalk.

In addition to uncovering striatal microglial signatures of traumatic stress and separately, voluntary wheel running, we were also interested in exploring potential overlap in signatures of running and stress on microglia. We found that stress-DEGs in rats with running wheel access did not significantly overlap stress-DEGs in rats without running wheel access (**Supp. Fig. 5C**); however, we identified a suite of eight genes (*Apoe, C1qa, Cst3, Grm3, Mt.co1, Pde4b, Rpl19,* and *Tmem176b*) that was differentially expressed in response to both stress and exercise. Notably, these genes are almost all involved in microglial activation (**Supp. Fig. 6**). Based on these data, we hypothesize that microglia are generally activated but exhibit different subtypes of responses to stress depending on exercise enrichment; however, future studies are needed to test this hypothesis.

We investigated microglial metabolism, pro-inflammatory activation (M1), and anti-inflammatory activation (M2) by tracking established transcriptional markers of these functional states. When all four experimental groups were analyzed together in a 2×2 design, microglia clustered into three subclusters, the exercise enrichment subcluster (more abundant in runners regardless of stress condition), relatively mature subcluster (more abundant in stressed versus non-stressed rats), and relatively immature subcluster. The exercise enrichment subcluster showed an increased M2 score compared to the relatively mature subcluster (**Fig. 5E**), further supporting distinct microglial signatures of stress and exercise, and supporting the idea that exercise promotes a shift toward anti-inflammatory activation. However, future research should validate the role of microglia activation, specifically the anti-inflammatory activation, in stress-induced decreases in running behavior.

In conclusion, our work demonstrates that a single traumatic stressor decreases voluntary wheel running and induces signatures of striatal microglial activation six weeks later. Further, we establish that striatal microglia display distinct molecular signatures associated with acute stress, exercise enrichment, and long-term running distances. Notably, the observed signature of microglia activation long after an acute stressor, coupled with the emergence of a unique exercise-associated subcluster displaying signatures of activation (M2) and phagocytosis, underscores the complex and multidimensional nature of microglial function. Both the unique functional profile of the microglia long after stress along with the exercise enrichment subcluster further support that microglial functional diversity extends beyond traditional M1 and M2 categories. The specific signature of changes in activation six weeks post-stressor also highlights the temporal complexities of microglial responses and may reflect unique forms of long-term priming, though future studies are needed to determine if priming has taken place. Future research should also investigate microglial functional diversity and causally manipulate striatal microglia to determine their functional role in modulating neural circuits and motivational deficits associated with stress.

## CRediT Author Contributions

**Meghan G. Connolly:** Conceptualization, Methodology, Formal Analysis, Software, Investigation, Writing – Original Draft, Writing – Review & Editing, Visualization, Project Administration. **Zachary V. Johnson:** Methodology, Formal Analysis, Software, Investigation, Data Curation, Writing – Review & Editing. **Lynna Chu:** Methodology, Formal Analysis, Software, Investigation, Data Curation, Writing – Review & Editing. **Nicholas D. Johnson:** Methodology, Formal Analysis, Software, Writing – Review & Editing. **Trevor J. Buhr:** Investigation. **Elizabeth M. McNeill:** Resources, Methodology. **Peter J. Clark:** Conceptualization, Investigation, Methodology, Resources, Writing – Review & Editing, Funding Acquisition. **Justin S. Rhodes:** Conceptualization, Methodology, Formal Analysis, Software, Investigation, Writing – Original Draft, Writing – Review & Editing, Supervision.

## Supporting information

Supplementary Results

