## Supplementary Figures for "Single-nucleus RNA sequencing reveals enduring signatures of acute stress and chronic exercise in striatal microglia"

### Slide 1
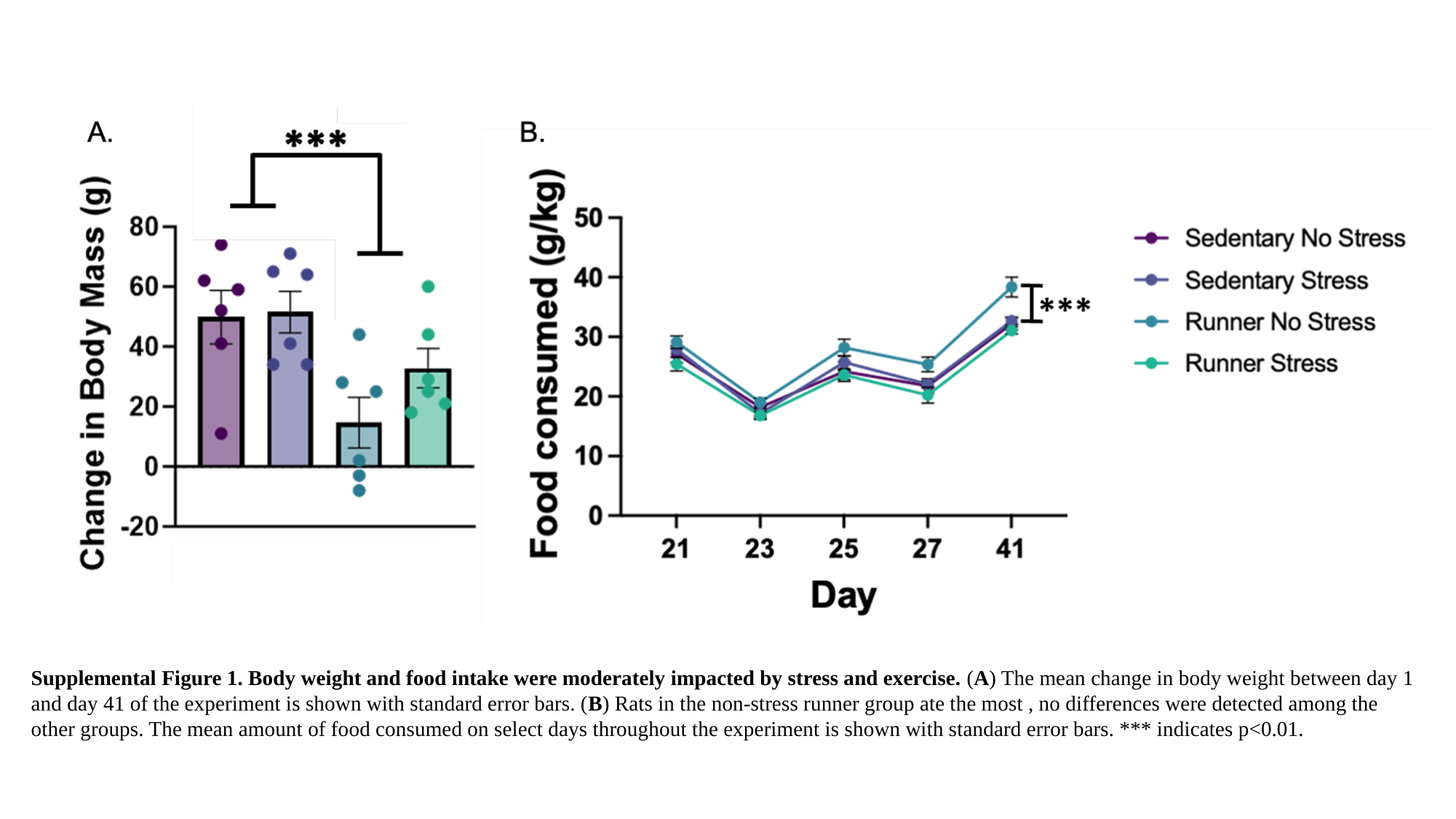

Supplemental Figure 1. Body weight and food intake were moderately impacted by stress and exercise. (A) The mean change in body weight between day 1 and day 41 of the experiment is shown with standard error bars. (B) Rats in the non-stress runner group ate the most , no differences were detected among the other groups. The mean amount of food consumed on select days throughout the experiment is shown with standard error bars. *** indicates p<0.01.

### Slide 2
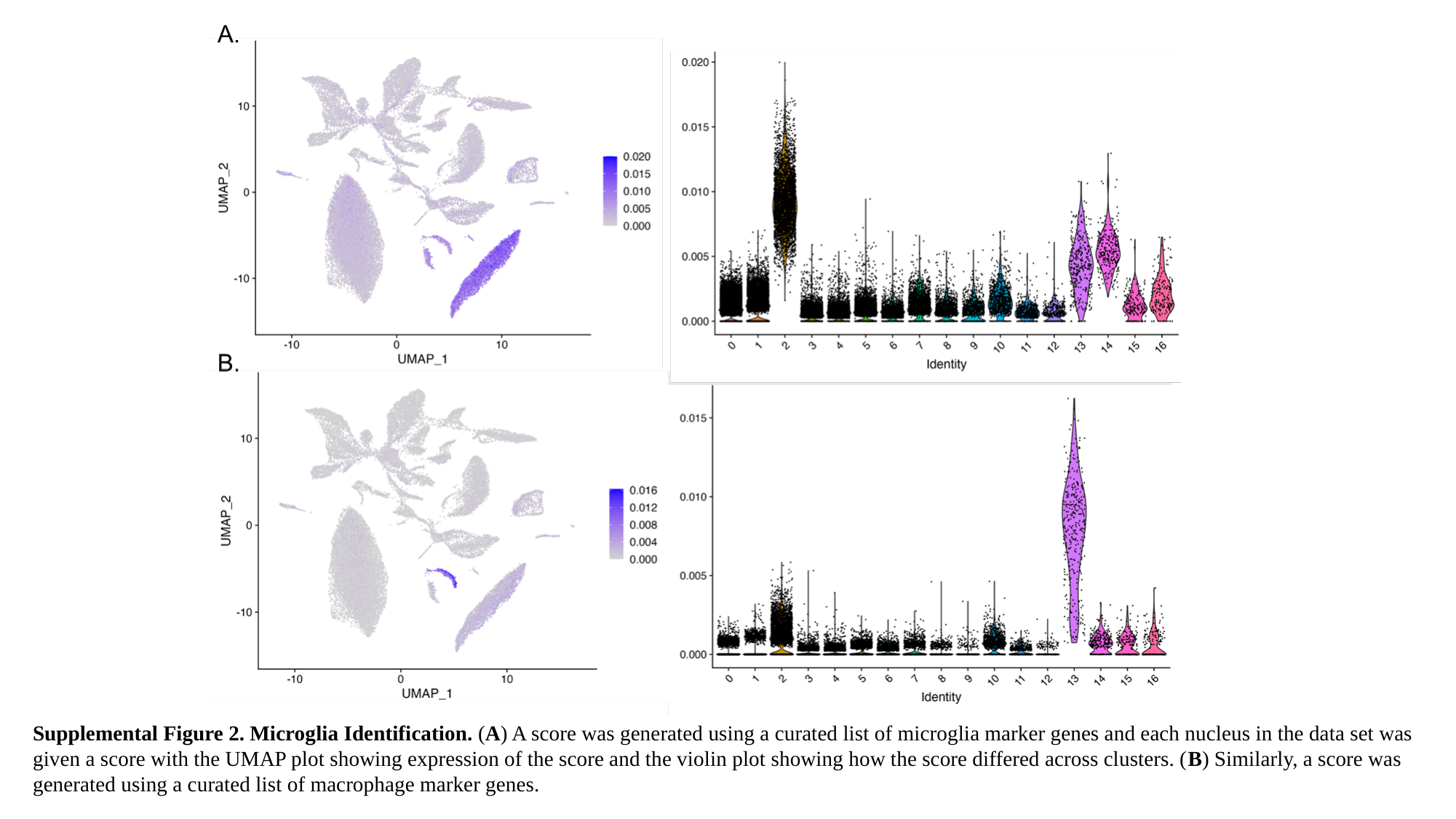

Supplemental Figure 2. Microglia Identification. (A) A score was generated using a curated list of microglia marker genes and each nucleus in the data set was given a score with the UMAP plot showing expression of the score and the violin plot showing how the score differed across clusters. (B) Similarly, a score was generated using a curated list of macrophage marker genes.

### Slide 3
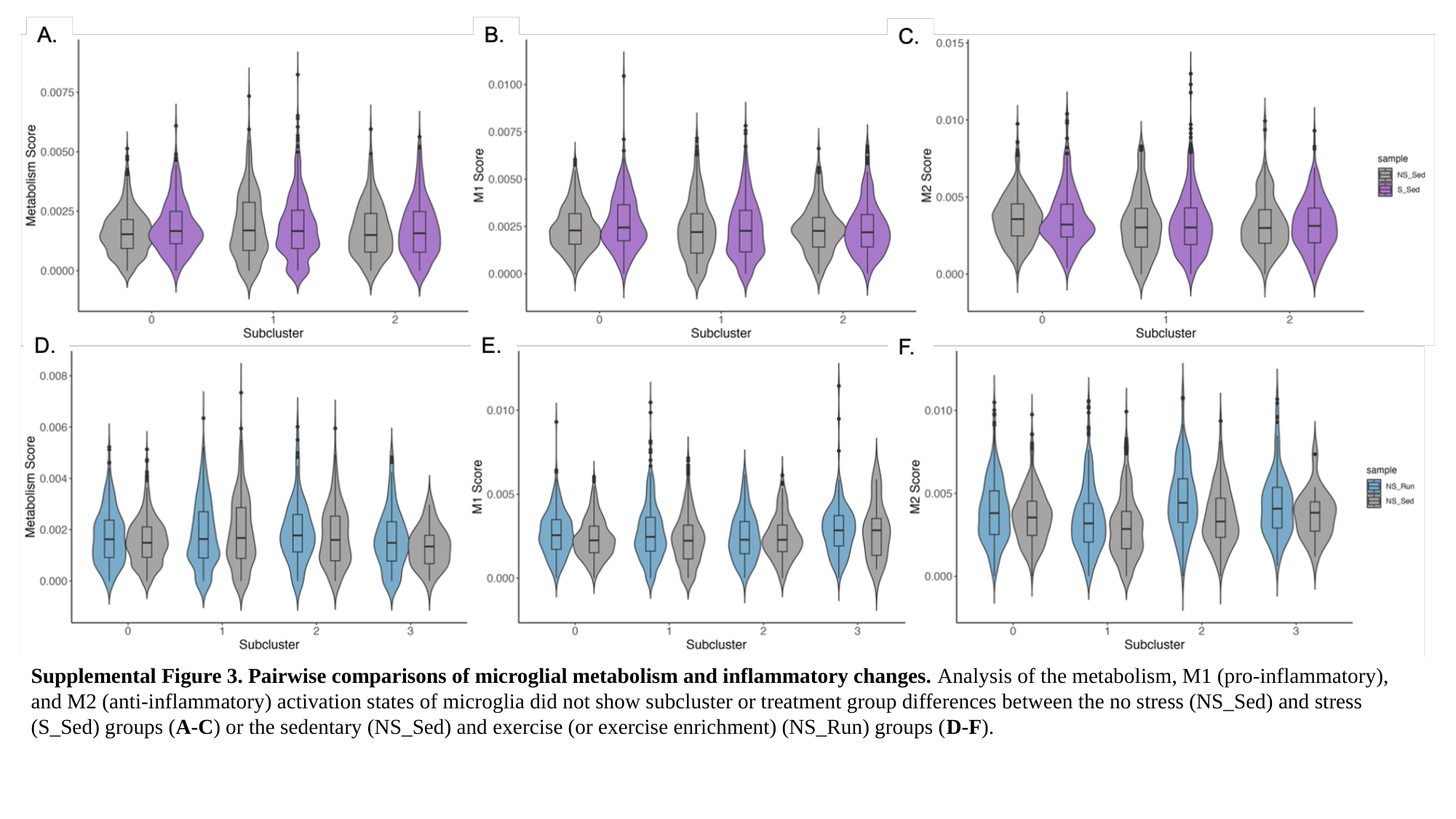

Supplemental Figure 3. Pairwise comparisons of microglial metabolism and inflammatory changes. Analysis of the metabolism, M1 (pro-inflammatory), and M2 (anti-inflammatory) activation states of microglia did not show subcluster or treatment group differences between the no stress (NS_Sed) and stress (S_Sed) groups (A-C) or the sedentary (NS_Sed) and exercise (or exercise enrichment) (NS_Run) groups (D-F).

### Slide 4
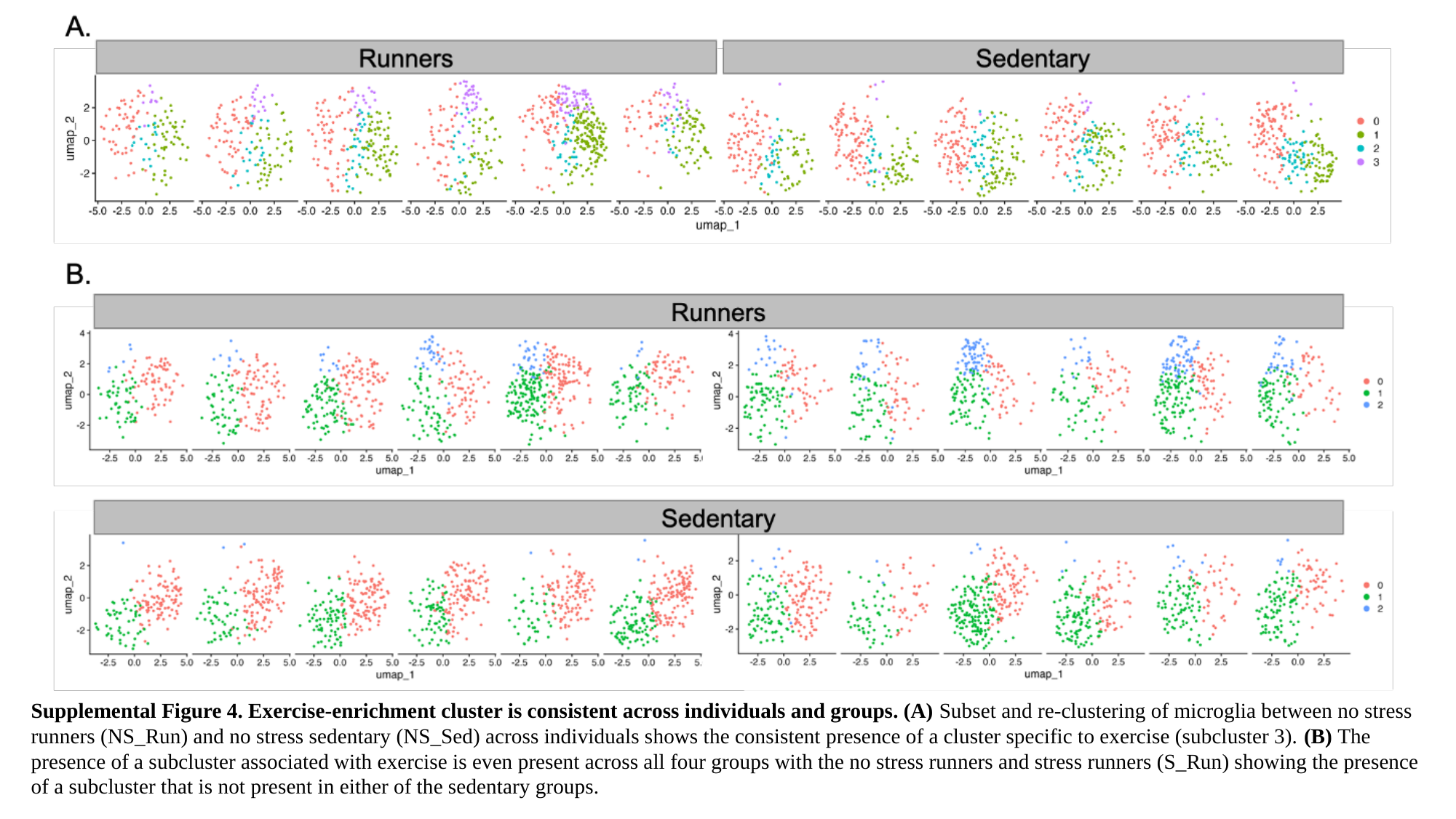

Supplemental Figure 4. Exercise-enrichment cluster is consistent across individuals and groups. (A) Subset and re-clustering of microglia between no stress runners (NS_Run) and no stress sedentary (NS_Sed) across individuals shows the consistent presence of a cluster specific to exercise (subcluster 3). (B) The presence of a subcluster associated with exercise is even present across all four groups with the no stress runners and stress runners (S_Run) showing the presence of a subcluster that is not present in either of the sedentary groups.

### Slide 5
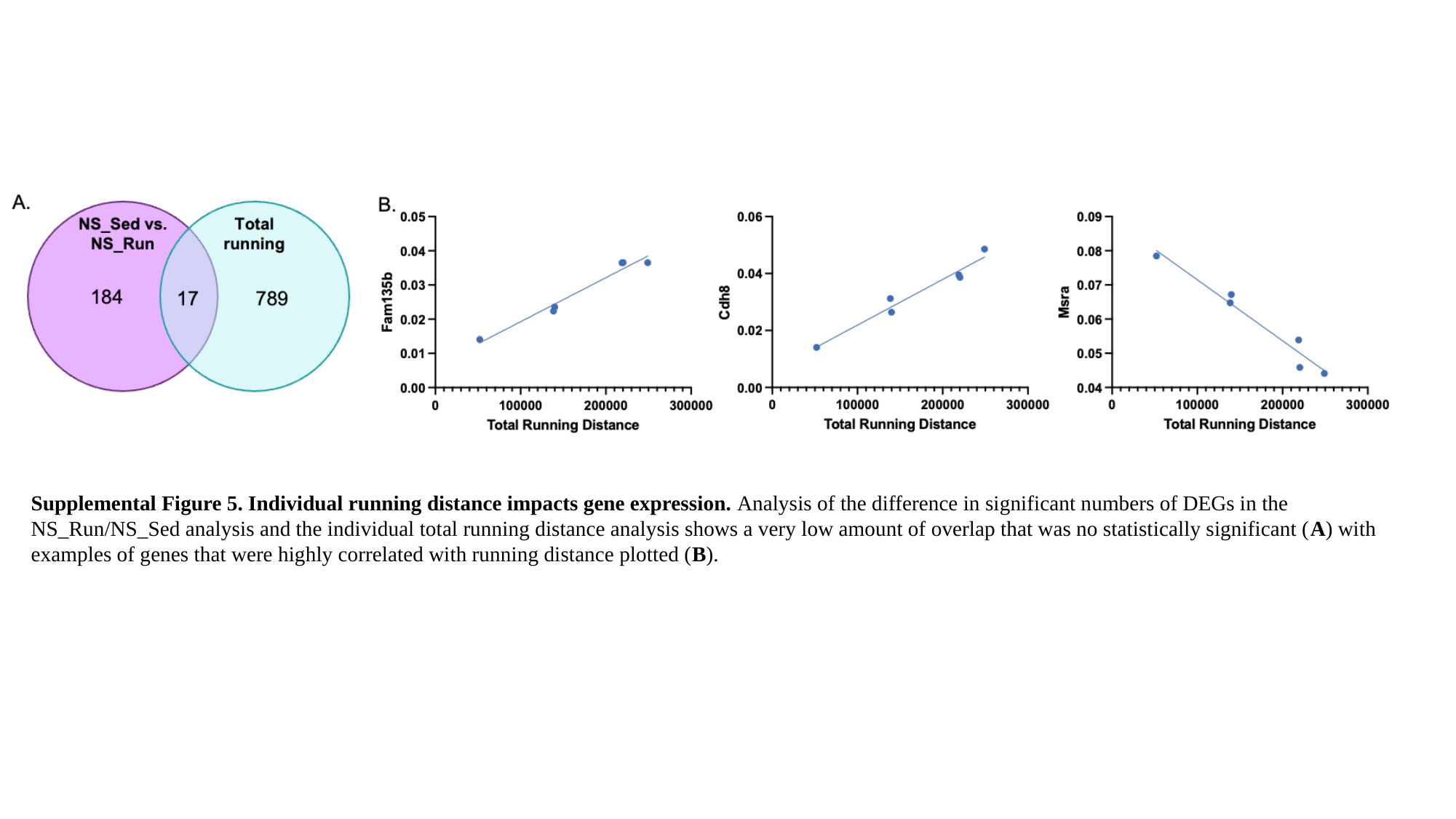

Supplemental Figure 5. Individual running distance impacts gene expression. Analysis of the difference in significant numbers of DEGs in the NS_Run/NS_Sed analysis and the individual total running distance analysis shows a very low amount of overlap that was no statistically significant (A) with examples of genes that were highly correlated with running distance plotted (B).

### Slide 6
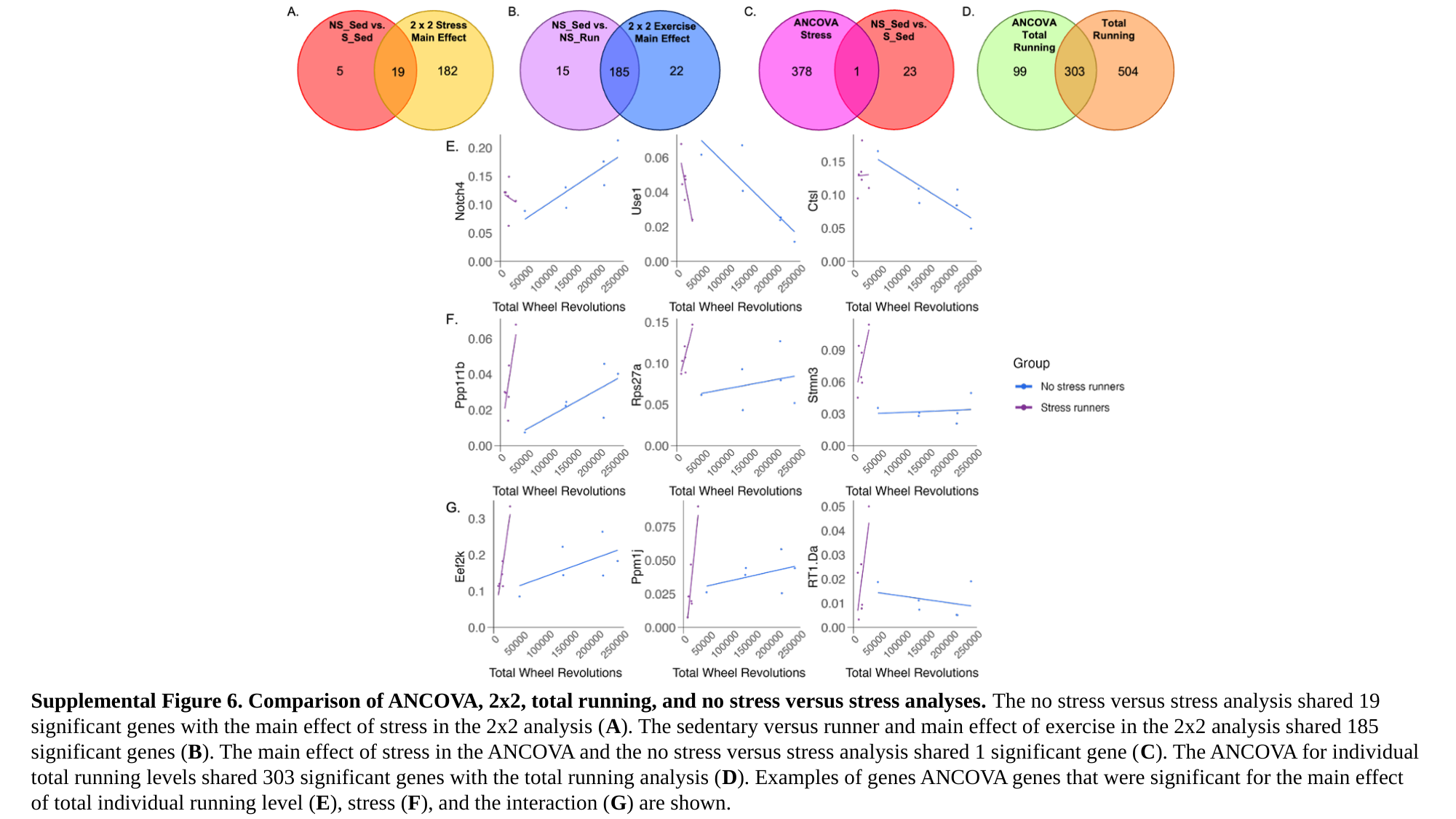

Supplemental Figure 6. Comparison of ANCOVA, 2x2, total running, and no stress versus stress analyses. The no stress versus stress analysis shared 19 significant genes with the main effect of stress in the 2x2 analysis (A). The sedentary versus runner and main effect of exercise in the 2x2 analysis shared 185 significant genes (B). The main effect of stress in the ANCOVA and the no stress versus stress analysis shared 1 significant gene (C). The ANCOVA for individual total running levels shared 303 significant genes with the total running analysis (D). Examples of genes ANCOVA genes that were significant for the main effect of total individual running level (E), stress (F), and the interaction (G) are shown.

### Slide 7
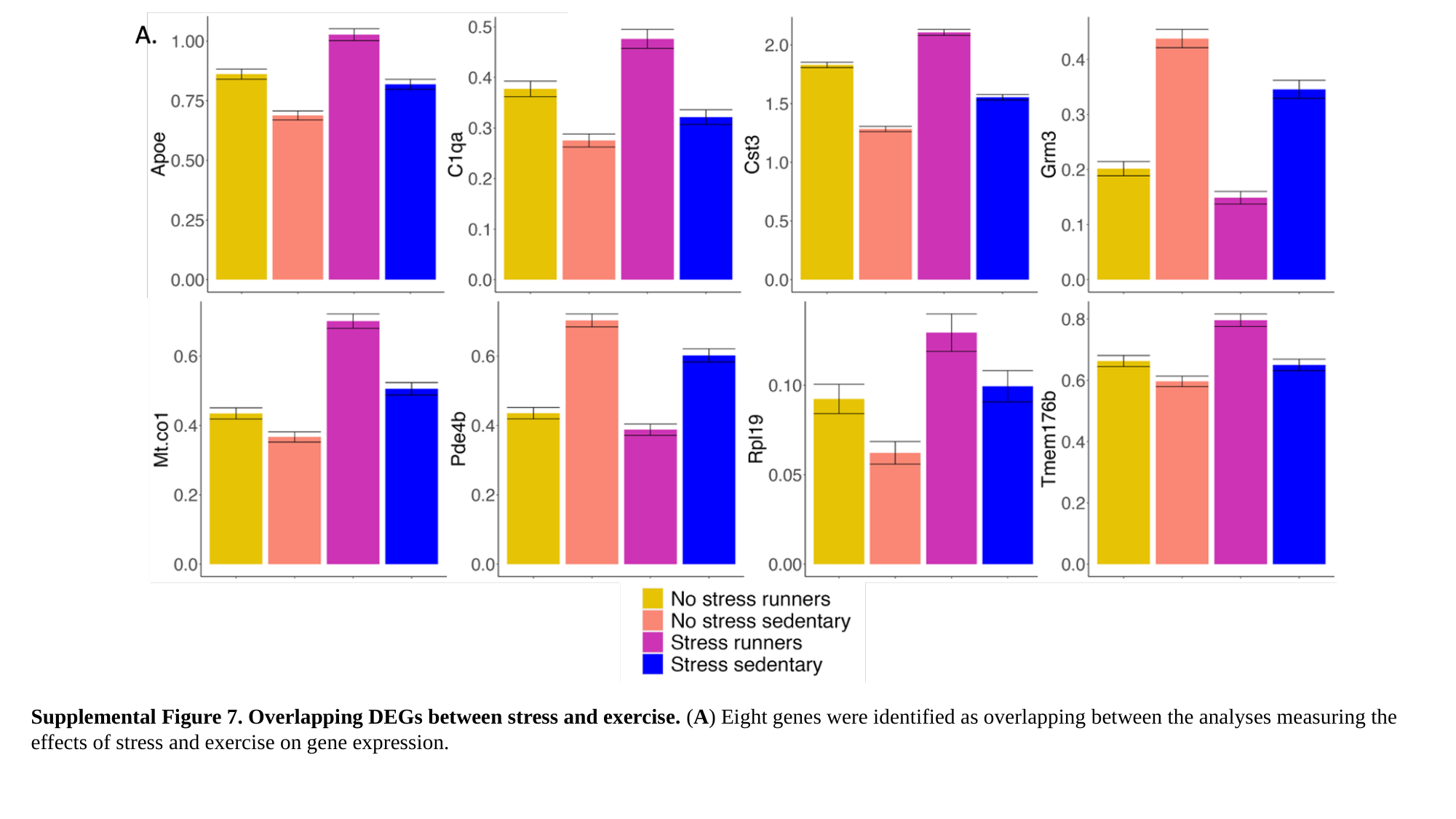

Supplemental Figure 7. Overlapping DEGs between stress and exercise. (A) Eight genes were identified as overlapping between the analyses measuring the effects of stress and exercise on gene expression.

### Slide 8
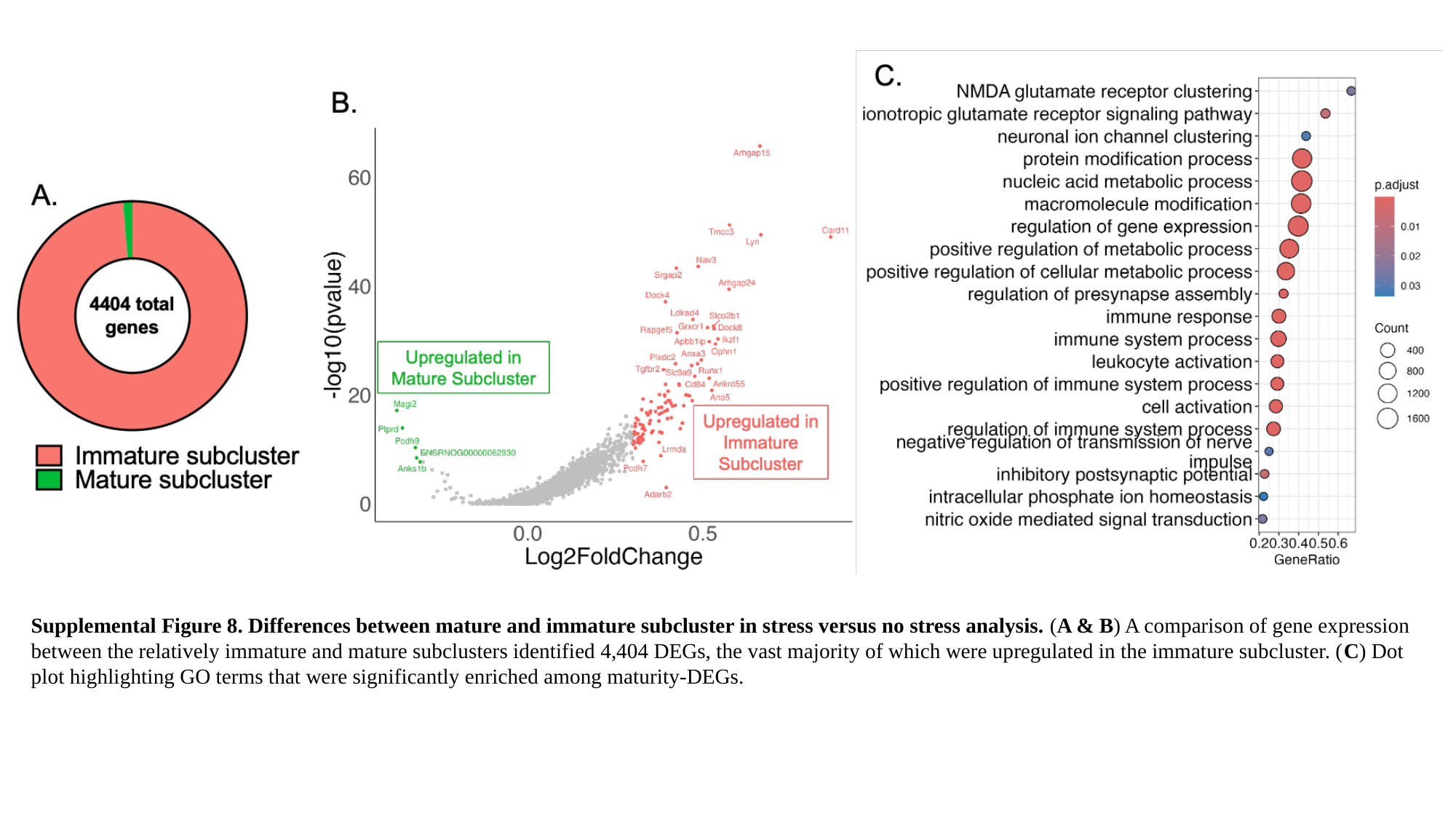

Supplemental Figure 8. Differences between mature and immature subcluster in stress versus no stress analysis. (A & B) A comparison of gene expression between the relatively immature and mature subclusters identified 4,404 DEGs, the vast majority of which were upregulated in the immature subcluster. (C) Dot plot highlighting GO terms that were significantly enriched among maturity-DEGs.

### Slide 9
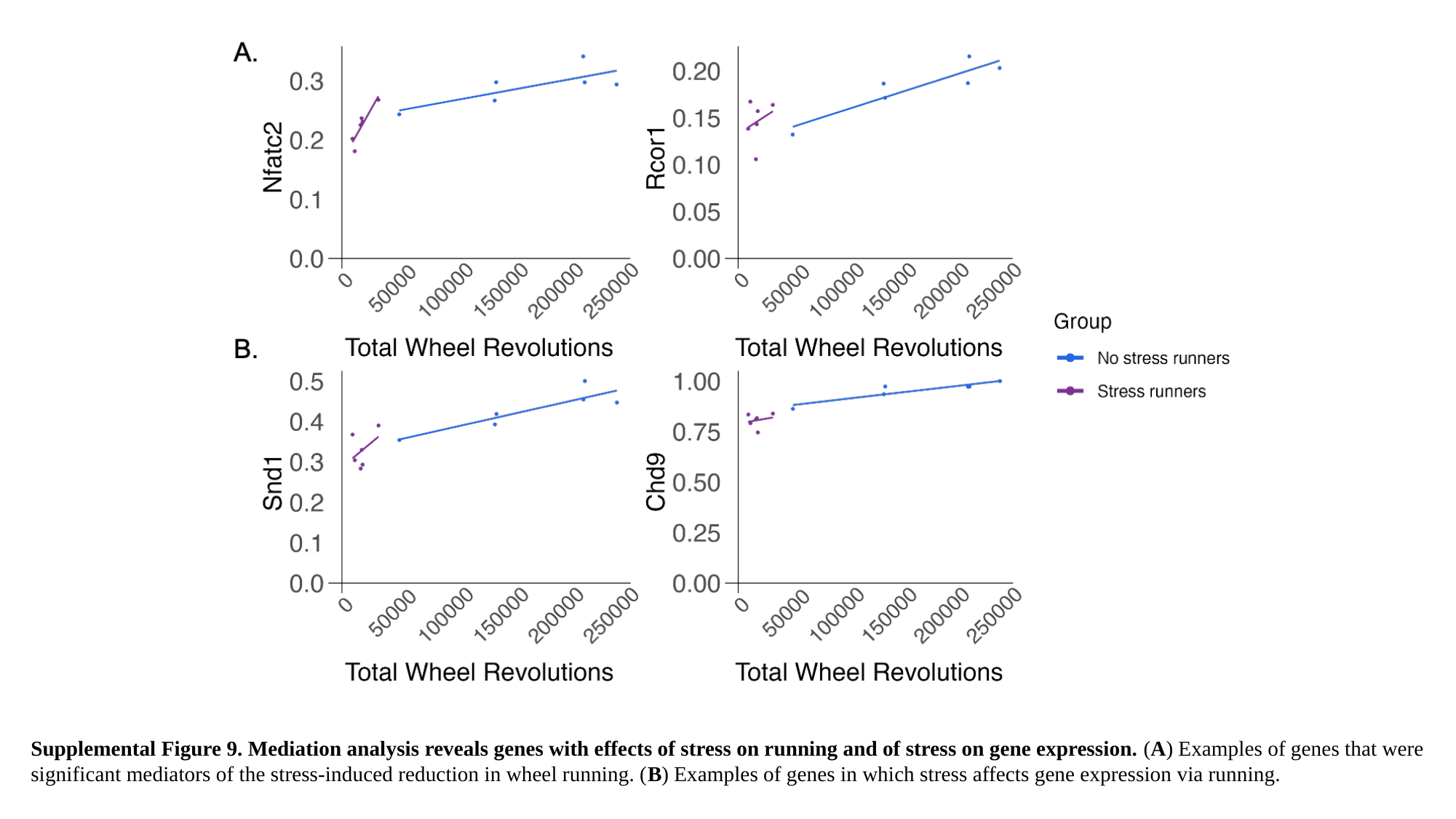

Supplemental Figure 9. Mediation analysis reveals genes with effects of stress on running and of stress on gene expression. (A) Examples of genes that were significant mediators of the stress-induced reduction in wheel running. (B) Examples of genes in which stress affects gene expression via running.
