## Supplementary Results for "Single-nucleus RNA sequencing reveals enduring signatures of acute stress and chronic exercise in striatal microglia"

**Rat Food Consumption and Body Weight**

Rats with access to running wheels gained less weight over the course of the study than sedentary rats (**Supp. Fig. 1A**). This was indicated by a significant main effect of access to a running wheel on weight gain (F(1, 20)=11.9, p=0.003). Rats in the No Stress Runner group ate more food than the other groups (**Supp. Fig. 1B**). This was indicated by a significant effect of day (F(4, 92)=158.0, p<0.0001), stress (F(1, 20)=8.0, p=0.01) and stress-by-exercise interaction (F(1, 20)=11.5, p=0.003). Posthoc analysis indicated the No Stress Runner group ate more than all other groups (all p<0.05), which ate similar amounts as each other.

**Mature versus Immature Subcluster in Stress versus No Stress Analysis**

To further characterize microglial subclusters, differential gene expression was analyzed in the mature versus immature subclusters. A Wilcoxon Rank Sum test identified 4,404 genes that exhibited differential expression between the mature versus immature subclusters (maturation-DEGs). The vast majority (4,346/4,404) of maturation-DEGs were upregulated in the immature subcluster compared to the mature subcluster (Supp. Fig. 8A & 8B) (Supp. Table 7). To investigate the functional significance of maturation-DEGs, we tested maturation-DEGs for enrichment of specific gene ontology (GO) terms (Supp. Fig. 8C) (Supp. Table 19). Maturation-DEGs were enriched for a large number of categories related to metabolism (e.g., Pik3r1, Nfkb1), immune system processes (e.g., Cd84, Il15), glutamate signaling (e.g., Grin1, Grid1) and synaptic changes (e.g., Nlgn1, Snca). The results from gene set enrichment analysis as well as differential gene expression analysis are consistent with altered glutamatergic signaling and enhanced metabolic activity in immature microglia relative to mature microglia.

**Alternative Bulk-seq Analysis of Differential Gene Expression**

In the main paper we analyzed gene expression differences between treatment groups (e.g., exercise, stress) using a mixed effects (glmer) negative binomial model with an offset term to adjust for library sizes among nuclei and dispersion estimated using a Gamma-Poisson model. A random effect was entered to estimate variation among individuals. The response was the total number of transcripts observed in the nucleus, hence the model estimates individual-level variance and nucleus-level variance (error term). In approximately 70% of the analyses, we found that the individual variance component was not estimable (results were “singular”). There is no universally accepted standard on how to deal with singularity in a model. One possibility is that the model is overfit. Other possibilities include numerical approximation or convergence issues. Thus, to evaluate the robustness of our results, we also analyzed the data a different way, using the SCtransform (Variance Stabilizing Transformations for Single Nuclei UMI Data) function in Seurat to process the data. This normalizes transcript count data to adjust for library sizes within nuclei. Further, we collapsed the data across all the nuclei in an individual effectively analyzing the data as if it were bulk-seq instead of single nuclei. We then analyzed the individual data points as a function of group using simple t-test, 2x2 ANOVA, linear regression using distance run as a covariate, and ANCOVA. P-values instead of q-values were used for SCtransform analyses to identify DEGs because the much smaller sample sizes in the SCtransform analyses (where the observational unit was the individual, rather than nuclei) resulted in a reduction in power. We found strong concordance between lists of DEGs from the SCtransform using p<0.05 and glmer using q<0.05. Because these are arbitrary cut offs, and do not match between analyses, we also compared concordance for the top 50 DEGs from each list. We ran the overlap in DEG lists between the two analyses for: Stress vs. No stress (all sedentary), Runner vs. No Runner (no stress), 2X2, simple linear regression, and ANCOVA. We ran hypergeometric tests to determine whether the overlap in DEG lists was significant between SCtransform and glmer analyses. For all pairs of analyses, the overlap in gene lists was significant (p<0.001). The significant overlap strongly supports the robustness of the results as presented in the main body of the paper.
